# Enhancer Tuning by Sequence-Specific Repressors

**DOI:** 10.64898/2026.09.11.751074

**Authors:** Lucia Ichino, Kaelan J. Brennan, Benjamin J. Mallory, Tomek Swigut, Stephanie C. Bohaczuk, Jaaved Mohammed, Devaraja G. Mudeppa, Saman Tabatabaee, Chunyang Ni, Andrew B. Stergachis, Joanna Wysocka

## Abstract

Enhancers integrate combinatorial inputs from sequence-specific transcription factors (TFs) and their activity must be calibrated to achieve precise spatiotemporal control of transcript dosage. Here we demonstrate that the sequence-specific repressors SNAI1 and SNAI2 (*i.e.* SNAIL and SLUG) quantitatively tune enhancer activity. In human neural crest cells, SNAI1/2 occupy a subset of active enhancers, where their depletion increases H3K27ac, chromatin accessibility, and enhancer regulatory potential. Changes in SNAI1/2 binding motifs contribute to enhancer divergence between human and chimpanzee, suggesting an evolutionary role for the repressor-mediated tuning. Single-molecule chromatin profiling using Deaminase-Assisted Fiber-seq (DAF-seq) reveals that individual enhancers toggle between an ensemble of open and nucleosome-dense chromatin states. While transcriptional activators increase the fraction of the open states, SNAI1/2 shift the equilibrium toward nucleosome-occupied states. This impedes binding of activator TFs, without fully repressing the enhancer. We propose that SNAI1/2 function as a molecular dimmer switch—modulating nucleosome dynamics to calibrate enhancer output.

## Introduction

Transcription of developmentally regulated genes is governed by transcription factors (TFs) that bind specific DNA sequence motifs within discrete regulatory elements termed enhancers.^1,2^ The interplay between expressed TFs and the arrangement and nature of their binding motifs determines the strength with which an enhancer is "activated" to increase expression of its target gene.^2,3^ Most research on enhancer activity regulation, especially in mammalian cells, has focused on activator TFs, which are critical for establishing and maintaining open chromatin states at enhancers and for enabling gene expression.^4–7^ Intriguingly, however, recent studies utilizing massively parallel reporter assays (MPRAs) or deep learning models of enhancer chromatin accessibility showed that DNA motifs recognized by sequence-specific repressors (*i.e.* transcription factors binding specific DNA sequences to mediate repression) are abundant at native enhancers and correlated with, respectively, weaker enhancer activation potential or predicted accessibility.^8–11^ Similarly, our previous analyses of evolutionary enhancer divergence revealed that presence of sequence-specific repressor motifs is correlated with enhancer activity decreases during evolution.^12^ Together, these observations suggest an underappreciated, yet potentially widespread role of sequence-specific repressor TFs in regulating enhancer activity, but the experimental evidence for such a role is still lacking.

SNAI1 and SNAI2 (hereafter SNAI1/2, also known as SNAIL and SLUG) are paralogous zinc-finger repressor TFs with broad roles in human development and cancer.^13,14^ They are most highly expressed in embryonic mesenchymal cell populations, such as the primitive streak, extraembryonic and embryonic mesoderm and neural crest, and their developmental role can be hijacked by cancer cells to promote invasion.^15–17^ Mechanistically, mammalian SNAI1/2 are best characterized as promoter-binding TFs, where they recognize E-boxes (CAGGTG) and recruit corepressors to silence expression.^14,18–22^ While the repressor function of SNAI1/2 is broadly recognized, in some contexts, they have also been shown to potentiate transcription.^23–25^ The repressive role of Snail, the fly ortholog of SNAI1/2, has long been studied in the context of *Drosophila* gastrulation and mesoderm formation.^14,26,27^ Snail directly represses genes that would otherwise specify ectodermal programs within the presumptive mesoderm, while the activator TF, Twist, mediates activation of mesodermal genes.^26,28,29^ In this system, Snail was proposed to act as a ‘short-range repressor’, capable of quenching activator binding within 50– 500 bp.^30–33^ When deployed at a promoter, Snail represses transcription, but when bound at an enhancer, it locally silences the target enhancer without preventing neighboring enhancers from regulating the same promoter.^30,31^ *Drosophila* studies therefore establish a precedent for Snail role in enhancer regulation, but in this context, Snail function appears dominant rather than modulatory – its cognate cis-regulatory element, whether enhancer or promoter, is actively silenced.^30,31^

To revisit the role of SNAI1/2 at enhancers in mammals, we turn to the neural crest, an embryonic cell population in which SNAI1/2 and TWIST1 are highly expressed and functionally important.^34–39^ Using *in vitro*-derived cranial neural crest cells (CNCCs) we show that SNAI1/2 occupy a subset of active enhancers, many of which are also bound by TWIST1. By leveraging human-chimp tetraploid cells,^40^ we further demonstrate that within shared human-chimp trans-regulatory environment, gains of SNAI1/2 motifs during evolution are associated with decreased enhancer acetylation. However, rather than silencing target enhancers, SNAI1/2 tune their activity by quantitatively dimming regulatory output, as measured by H3K27ac, chromatin accessibility, activator TF binding and ability to drive reporter gene expression. We thus refer to these as “SNAI1/2-tuned enhancers”. To investigate the mechanism of enhancer tuning, we deploy single-molecule approaches, Fiber-seq and DAF-seq. These studies reveal remarkable heterogeneity of nucleosome patterns at active enhancers within a population of cells existing in the same defined cellular state. Depletion of TWIST1 or SNAI1/2 does not reduce this diversity of enhancer chromatin states; instead, the two factors shift the distribution in opposite directions—TWIST1 toward open, nucleosome-depleted states and SNAI1/2 toward closed, nucleosome-dense states. Together, these findings identify enhancer tuning through rebalancing of chromatin states as a mechanism by which repressive TFs calibrate regulatory activity — and provide a route through which DNA sequence variation can quantitatively modulate enhancer output over evolutionary time.

## Results

### Cross-species comparison and deep learning suggest a role for SNAI1/2 motifs in negative modulation of enhancer activity

Analyzing DNA sequence features that underlie quantitative divergence in enhancer acetylation between closely related species, such as humans and chimpanzees, can reveal the transcription factors that regulate enhancer activity.^12,41^ We hypothesized that if sequence-specific repressors such as SNAI1/2 negatively modulate enhancer activity, species-specific sequence changes that weaken SNAI1/2 DNA binding should lead to increased enhancer H3K27ac (a chromatin mark correlated with enhancer activity), and conversely, those that strengthen the binding should result in decreased H3K27ac. To test this, we turned to human/chimpanzee tetraploid hybrid induced pluripotent stem cells (iPSCs) that were recently generated by cell fusion to investigate regulatory divergence.^40,42,43^ This system offers a unique opportunity to study the effects of cis-acting DNA sequence variation within a constant *trans* environment, composed of proteins and RNAs encoded by both the human and chimpanzee genomes.

To enable allele-resolved analyses, we first performed PacBio long-read sequencing to generate haplotype-phased assemblies of the tetraploid genome, allowing sequencing reads to be accurately assigned to their human or chimpanzee haplotype of origin. We then differentiated the hybrid iPSCs to cranial neural crest cells (CNCCs) using our established protocol,^12,39,44,45^ and performed chromatin immunoprecipitation sequencing (ChIP-seq) for SNAI2 and H3K27ac, mapping reads to the phased genomes to obtain allele-specific signal readout (Figure 1A, Table S1). We then mapped the well-characterized SNAI1/2 E-box binding motif — determined *in vitro* by SELEX^46^ — onto the human and chimpanzee resolved haplotypes (Figure 1A). This approach allowed us to identify DNA sequence variants between the human and chimpanzee genomes that alter predicted motif strength, and to directly measure their effects on SNAI2 occupancy and H3K27ac levels at the same distal elements.

**Figure 1:**
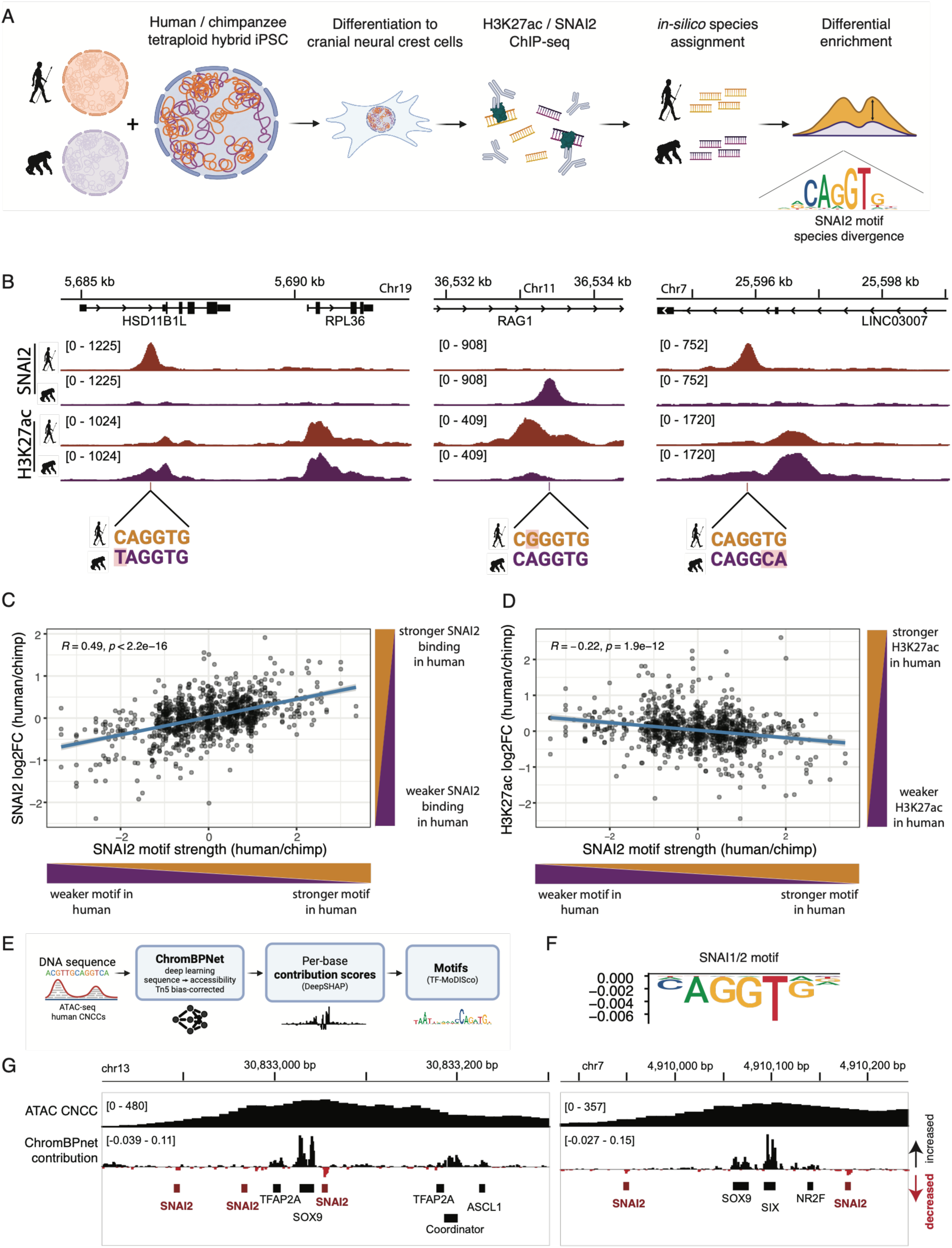
Cross-species divergence and deep-learning models implicate SNAI1/2 motifs in negative regulation of H3K27ac and ATAC-seq at enhancers. (A) Schematic representation of workflow for human/chimpanzee tetraploid hybrid cells ChIP-seq analysis (created with BioRender.com). (B) ChIP-seq tracks from human/chimpanzee hybrid line, split by assigned species, showing enhancers with mutations in the SNAI1/2 binding motif (mutations are shaded in red). (C,D) Scatterplots showing distal regulatory elements overlapping SNAI2 ChIP-seq peaks, plotting correlations between changes in ChIP-seq enrichment in human vs chimp reads and changes in SNAI2 motif^46^ FIMO^80^-calculated p-value (-log10(SNAI2 motif p-value human / SNAI2 motif p-value chimp)). If multiple motifs mapped to a single enhancer, the mean of p-value ratios was calculated. Pearson R and p-value are shown. E) Schematic of ChromBPNet analysis (created with BioRender.com). F) The SNAI2 motif negatively contributes to ChromBPNet’s prediction of chromatin accessibility in human CNCCs. The motif is shown as a contribution weight matrix (CWM), where the height of the base represents its importance for the prediction. G) Examples of enhancers where ChromBPNet model shows negative scores of contribution to counts at SNAI1/2 motifs.

These analyses revealed a reciprocal relationship: variants that perturb the SNAI1/2 motif reduce SNAI2 occupancy at the corresponding candidate enhancer, but this is accompanied by a gain in H3K27ac signal; conversely, stronger motifs are associated with higher SNAI2 binding and lower acetylation (Figures 1B-1D). This inverse relationship between SNAI1/2 motif strength/binding and H3K27ac levels across species suggests that SNAI2 occupancy negatively modulates enhancer activity, as approximated by H3K27ac. Importantly, this effect is quantitative – rather than resulting in enhancer decommissioning, gains of stronger SNAI1/2 motifs are associated with moderate, up to four-fold decreases in H3K27ac level (Figure 1D). We propose that such tuning mechanism could be exploited over evolutionary time to allow for precise modulation of enhancer activity. Consistent with a potential functional role of SNAI1/2 in enhancer regulation, SNAI1/2 motif occurrences within SNAI1/2 distal ChIP-seq peaks showed stronger evidence of purifying selection than unbound controls, both by cross-species constraint^47–49^ and by a joint estimate of constraint from between-species divergence and human polymorphism^50^ (Figure S1A and S1B).

Independent support for quantitative enhancer tuning by SNAI1/2 came from a deep learning model trained on chromatin accessibility data that learns to predict accessibility directly from DNA sequence (ChromBPNet).^51^ We trained ChromBPNet on ATAC-seq data from our human *in-vitro* derived CNCCs (Figure 1E). ChromBPNet model performance was high (∼0.7 Pearson R for counts performance) and consistent through cross-validation (Figures S1C and S1D1). Moreover, the same motif patterns for key human CNCC TFs were discovered across the cross-validation models, collectively indicating that our training approach was remarkably stable (Figure S1E). As expected, the model identified well-known cranial neural crest TFs — TWIST1, TFAP2A, NR2F1, SOX9, and SIX1/2 — and other well-known TFs like CTCF, as positive contributors to accessibility (Figures S1E and S1F). In contrast, the SNAI1/2 motif came up as a negative contributor, namely its presence was predicted to decrease accessibility (Figures 1F, 1G, S1E, and S1F), consistent with SNAI1/2 counteracting chromatin opening at the bound elements. We note that the negative contribution scores of SNAI1/2 to accessibility were typically smaller in magnitude than positive contribution scores of the strongest activators (Figures 1G and S1F), which again suggests tuning, rather than dominant repression of enhancer activity by SNAI1/2.

### SNAI1 and SNAI2 function redundantly in CNCCs and bind a subset of active enhancer elements

We next sought to mechanistically validate the binding and regulation of active enhancers by SNAI1/2. To this end, we employed a degron-tagging approach by editing human embryonic stem cells (ESCs) to fuse the endogenous SNAI1 or SNAI2 proteins with the dTAG-inducible FKBP12^F36V^ degron,^52,53^ the fluorophore mNeonGreen, and a V5 epitope tag (collectively referred to as the FNV tag) (Figure 2A).^41^ We also generated a SNAI1-FNV line in SNAI2 knockout (KO) background to assess potential redundancies between the two homologs, as previously suggested (Figure 2A).^54^ Following differentiation of these hESC lines into CNCCs, we verified stable expression of the tagged proteins and lack of expression in the KO (Figures S2A–S2D) and performed RNA-seq after acute (3 hours) or prolonged (1 and 3–4 days) protein depletion with dTAG. We also performed ChIP-seq for SNAI1 and SNAI2 using endogenous or V5 antibodies to compare genome-wide binding patterns.

**Figure 2:**
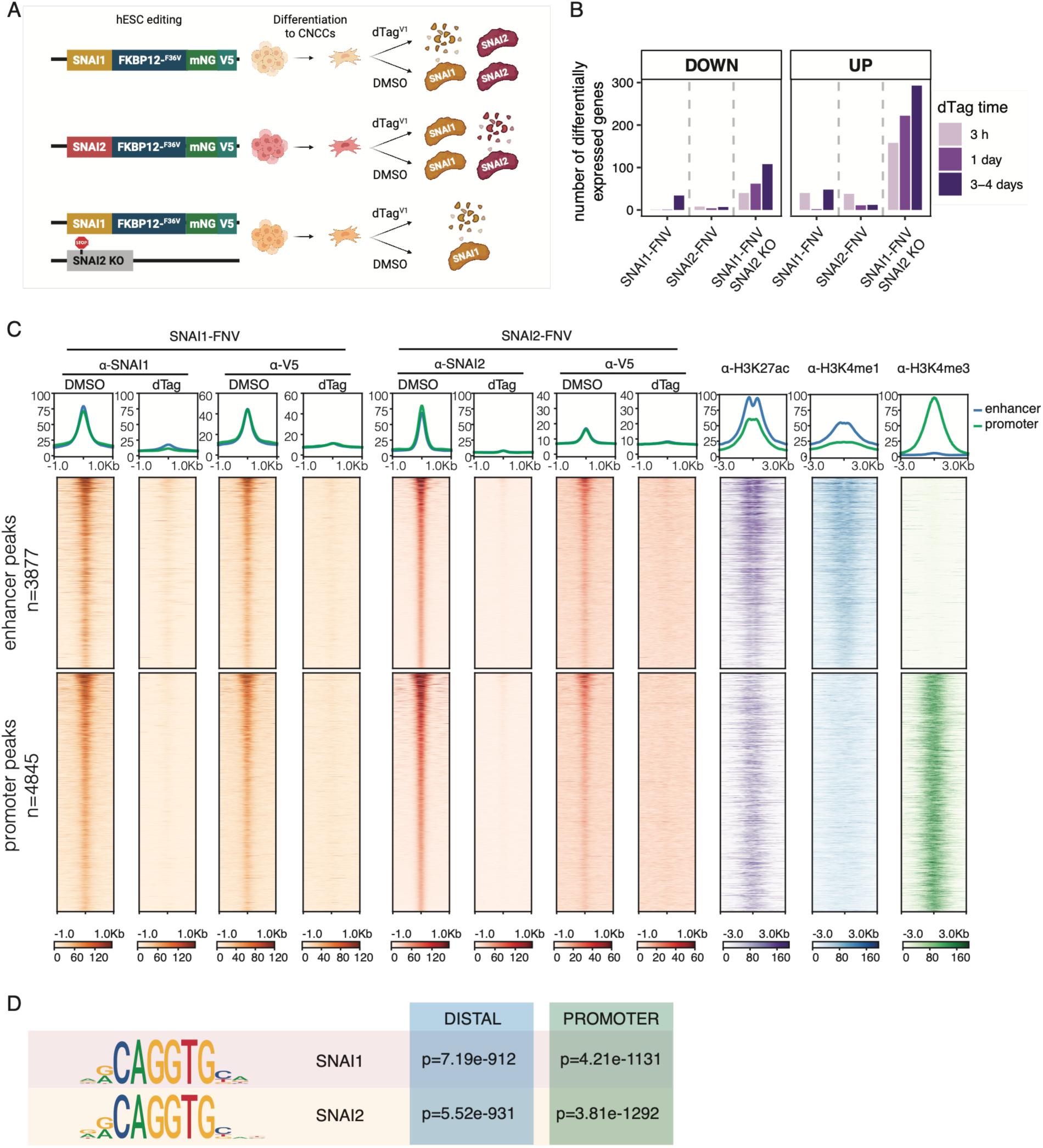
SNAI1 and SNAI2 act redundantly in CNCCs and occupy a subset of active enhancers. (A) Schematic representation of gene editing and protein depletion strategy (created with BioRender.com). Three lines were generated: SNAI1-FNV, SNAI2-FNV, SNAI1-FNV and SNAI2 KO. Two clones were used for each line. B) Numbers of differentially expressed genes obtained by RNA-seq in CNCCs, in the indicated lines and time points of dTAG treatment. (C) Heatmap of SNAI1 and SNAI2 ChIP-seq binding (in SNAI1-FNV SNAI2 KO or SNAI2-FNV lines respectively, 3 h DMSO/dTAG treatment) and histone marks (ChIP-seq in unedited H9 CNCCs) at promoter-proximal or distal SNAI1/2 peaks, ranked by anti-SNAI2 DMSO average signal (see Methods and Figure S2G for peaks annotation). Peaks annotated as “H3K27me3 enriched regions” are not shown in this heatmap. H3K4me3 data is from Prescott et al.^12^. Units: RPKM-normalized reads. (D) Enrichment of SNAI1 and SNAI2 motifs (HOCOMOCO)^81^ in promoter-proximal or distal SNAI1/2 peaks with p-values (MEME Analysis of Motif Enrichment).

RNA-seq analysis revealed functional redundancy between SNAI1 and SNAI2: depletion of either factor alone caused minimal transcriptome changes. In contrast, SNAI1 depletion in the SNAI2 KO background produced progressive transcriptional changes, with misregulated gene counts rising at longer depletion times (Figures 2B and S2E). This is consistent with mouse genetic studies demonstrating SNAI1/SNAI2 redundancy during chondrogenesis.^54^ The predominance of upregulated genes, together with their preferential overlap with SNAI1/2 ChIP peaks (Figure S2E), is consistent with the well-established roles of SNAI1 and SNAI2 as transcriptional repressors.^14^ Notably, transcriptional changes at target genes were already detectable after just 3 hours of depletion, indicating a rapid regulatory response to the SNAI1/2 loss (Figure 2B).

Analysis of ChIP-seq profiles identified 9,190 high-confidence peaks strongly enriched for SNAI1/2 binding motifs, at which signal was lost upon dTAG treatment (Figures 2C and 2D). SNAI1 and SNAI2 occupancy was well correlated across these sites, consistent with the functional redundancy observed by RNA-seq (Figure S2F). As expected, given their established role in promoter repression, many (n=4,845) of the SNAI1/2 peaks corresponded to promoters. However, a substantial fraction of the peaks (n=3,877) mapped to promoter-distal elements enriched for the enhancer marks H3K27ac and H3K4me1 and depleted of the promoter mark H3K4me3 (Figures 2C and S2G). Thus, SNAI1 and SNAI2 occupy a large number of active enhancer elements in CNCCs.

### SNAI1/2 negatively modulate enhancer output without complete repression

We next sought to determine whether SNAI1/2 have a functional impact at bound enhancers. Given the redundancy described above, we performed all subsequent experiments using the SNAI1-FNV line in SNAI2 knockout background. To evaluate the effect of SNAI1/2 at enhancers, we first performed H3K27ac ChIP-seq and ATAC-seq in CNCCs following acute or prolonged dTAG treatments. SNAI1/2 depletion increased H3K27ac and ATAC signals at over 1,000 genomic regions, whereas fewer regions showed decreased signals (Figure 3A). The upregulated regions were enriched for SNAI1/2 ChIP-seq binding (Figure 3B). In contrast, the downregulated regions showed limited overlap with SNAI1/2 ChIP-seq peaks and were most abundant at the long duration (3-day) dTAG time point, suggesting they likely represent indirect effects (Figure 3B). The upregulated and directly bound regions included both promoters and distal active enhancers, pointing to a direct effect of SNAI1/2 binding on enhancer attenuation, in addition to the well-established role in promoter regulation (Figures 3C and S2G).^14,18–22^ These distal, SNAI1/2-regulated elements were also devoid of repressive marks H3K27me3 and H3K9me3, further supporting the idea that under unperturbed conditions they are active (Figure S2H). Notably, we found that—perhaps counterintuitively—these distal regions displaying increased H3K27ac in short dTAG treatments tended to be among the enhancers with the highest H3K27ac levels in DMSO, suggesting that SNAI1/2 regulates strong enhancers (Figure 3D). Furthermore, H3K27ac and ATAC fold-changes were positively correlated at these distal, SNAI1/2 bound elements, supporting a mechanism that affects both readouts of enhancer function (Figure 3E). Nonetheless, regression slope (H3K27ac on ATAC) was progressively steeper with increasing time of dTAG treatment, as H3K27ac increased more slowly than chromatin accessibility after SNAI1/2 depletion (Figure 3E, Table S2), suggesting that accessibility changes may be a primary effect of SNAI1/2 loss.

**Figure 3:**
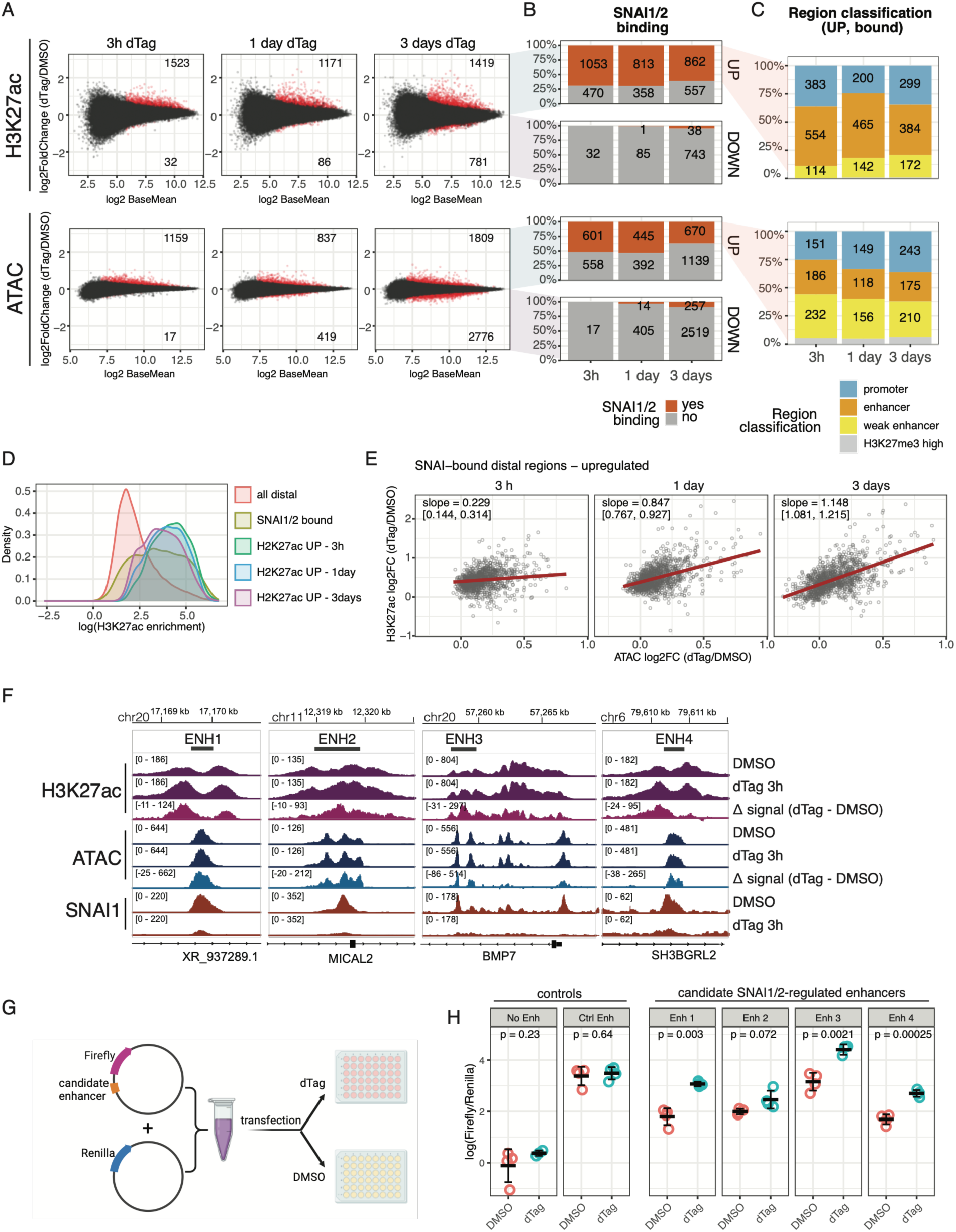
SNAI1/2 depletion quantitatively increases enhancer acetylation, accessibility and reporter output. (A) Scatterplots visualizing differential enrichment analysis for H3K27ac ChIP-seq and ATAC-seq in SNAI1-FNV SNAI2 KO line at different dTAG treatment time durations. Signal was calculated over 2 kb regions around all reproducible ATAC summits. Red indicates significantly changed regions (DEseq p-value< 0.05). (B) Fraction of significantly upregulated or downregulated ATAC regions that overlap a SNAI1/2 ChIP-seq peak. (C) Functional annotation of the upregulated and SNAI1/2 bound regions (see Figure S2G). (D) Density plot showing distribution of H3K27ac signal at different sets of regions including H3K27ac significantly upregulated regions from panel A. (E) Scatterplots comparing H3K27ac changes to ATAC-seq changes. Plotted points represent the union of regions that are (1) SNAI1/2-bound, (2) classified as "enhancer" or "other distal element," and (3) significantly upregulated in H3K27ac and/or ATAC-seq signal at one or more time points. Regression line and 95% confidence interval are estimated marginal effects from a linear mixed-effects model (H3K27ac log2FC ∼ ATAC log2FC × time point, with a random intercept per region) fit jointly across all three time points. (F) Genome browser tracks showing examples of SNAI1/2 tuned enhancers (SNAI1-FNV SNAI2 KO cell line). (G) Schematic representation of luciferase assay (created with BioRender.com). (H) Luciferase assay showing normalized firefly activity for different constructs and dTAG time points in SNAI1-FNV SNAI2 KO line. Four replicates per condition (individual wells of cells). Representative of at least 2 independent experiments. P-value: unpaired two-sample Welch’s *t*-test.

Overall, our results are in agreement with a model in which SNAI1/2 quantitatively tune the strength of active enhancers, without fully repressing or decommissioning them. However, because H3K27ac and ATAC-seq signals are only correlates of enhancer activity, we sought to measure enhancer activity directly using a dual luciferase reporter assay. We cloned a set of candidate SNAI1/2-regulated enhancers (Figure 3F) upstream of a minimal SV40 promoter driving Firefly luciferase expression and co-transfected these constructs with a Renilla control plasmid (Figure 3G). The tested elements behaved as active enhancers, showing Firefly activation relative to the promoter-only control (Figure 3H). Notably, SNAI1/2 depletion via dTAG increased enhancer output, supporting a role for SNAI1/2 in quantitative modulation, or “tuning”, of enhancer strength (Figure 3H). Finally, we analyzed signatures of evolutionary constraint, this time limiting the analysis to enhancers not only bound but also regulated by SNAI1/2. Strikingly, we found that evidence for purifying selection increased further at the SNAI1/2 motifs overlapping tuned enhancers (Figure S1A and S1B).

### SNAI1/2 interfere with activator recruitment at co-bound enhancers

Early studies in flies proposed that Snail, the *Drosophila* ortholog of SNAI1/2, interferes with transcriptional activators in a localized manner ("short-range repression"), but the supporting evidence was limited to several exemplary loci.^30,31^ To investigate the relationship between SNAI1/2 and activators in human CNCCs, we first leveraged ChromBPNet,^51^ which predicts ATAC-seq data directly from DNA sequence. We trained single-task models on ATAC-seq data collected from CNCCs in control and SNAI1/2-depleted conditions and interpreted the models to capture the sequence-encoded effects of SNAI1/2 depletion on chromatin accessibility (Figure S1). We selected candidate SNAI1/2-tuned enhancers at which the dTAG model predicted higher ATAC-seq signal than the DMSO model (Figure S3A) and asked how the per-base-pair contributions to accessibility across each enhancer changed in the dTAG model. The SNAI1/2 motif showed loss of negative contribution, which was accompanied by increased contribution scores at intervening activator motifs (Figure S3A). As further validation, we computationally mutated the SNAI1/2 motif. Mutating the motif increased predicted ATAC-seq signal under the DMSO model but produced minimal change under the dTAG model (Figures 4A and S3B), as expected if the motif is already non-functional once SNAI1/2 is depleted. In the DMSO model, mutating the SNAI1/2 motif led to loss of negative contribution at the mutated site, consistent with loss of SNAI1/2 binding, and increased contribution scores at all intervening activator motifs as well, suggesting that SNAI1/2 negatively modulates activators broadly across the enhancer (Figure 4A).

**Figure 4:**
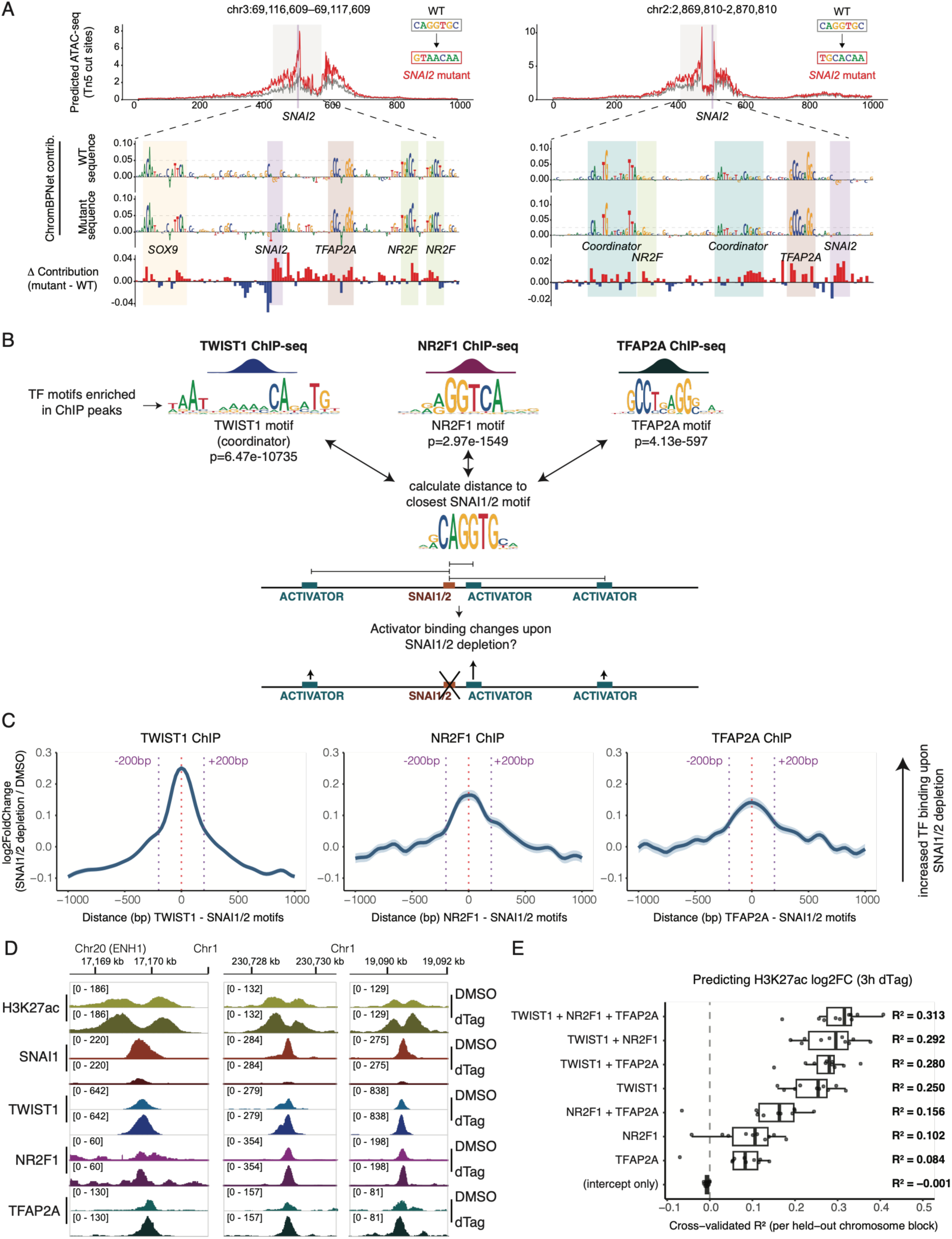
SNAI1/2 interfere with activator TF binding within ∼200 bp of their motif. (A) ChromBPNet predicts increased accessibility upon mutating SNAI2 motifs across enhancers. The trained DMSO ChromBPNet model was used to predict Tn5 cut site coverage across wildtype sequences (grey) and sequences in which the SNAI2 motif was mutated (red). The underlying contribution tracks show that as SNAI2 motifs become less negatively contributing, proximal motifs for activators tend to gain importance. The delta contribution track summarizes how each base across the sequence changes in contribution upon the mutation of the SNAI2 motif (red is increase in contribution, blue is loss of contribution). (B) Experimental workflow: TWIST1, NR2F1 and TFAP2A ChIP-seq was performed in SNAI1-FNV SNAI2-KO CNCCs (DMSO and 3h dTAG). The top motif enriched in each peak set is shown with enrichment p-value (MEME Analysis of Motif Enrichment). For each motif in ChIP peaks, we calculated the distance to the closest SNAI1/2 motif, and the log2 fold-change (dTAG / DMSO) of ChIP-seq enrichment at the overlapping TF peak. (C) Kernel-smoothed log2 fold-change of ChIP-seq enrichment for TWIST1, NR2F1, or TFAP2A at distal regulatory regions in SNAI1-FNV SNAI2-KO CNCCs, 3 h dTAG vs DMSO. For each peak, the summit was matched to the nearest TF-specific motif occurrence (≤ 100 bp), and its position was expressed as a signed distance to the closest SNAI1/2 motif, oriented relative to SNAI1/2 motif strand. A Gaussian kernel smoother (bandwidth = 100 bp) was applied across all signed distances to estimate the mean log2 fold-change as a function of relative motif spacing; shaded area indicates the weighted standard error of the kernel estimate. (D) Examples of genome browser tracks showing increased binding of activators upon SNAI1/2 depletion. Shown is ChIP-seq data of H3K27ac and TFs in SNAI1-FNV SNAI2-KO CNCCs (DMSO or 3 h dTAG). (E) Cross-validated predictive performance of linear models estimating H3K27ac log2 fold-change at distal regulatory regions bound by all three TFs upon SNAI1/2 depletion. The ChIP-seq log2 fold-changes of TWIST1, NR2F1, and TFAP2A were used as predictors, individually or in combination, alongside an intercept-only null model. Cross-validation was blocked by chromosome. Points show per-fold R², each computed against the mean of its own held-out block; boxes show median and interquartile range. The value at right of each row is the pooled out-of-fold R², obtained from the concatenated held-out predictions of all folds.

To validate these predictions experimentally, we performed ChIP-seq for TWIST1, NR2F1, and TFAP2A in SNAI1-FNV-SNAI2-KO CNCCs, with and without acute (3 hours) SNAI1/2 depletion. We first confirmed that the top enriched motif in each ChIP-seq dataset matched the expected binding motif for that TF (Figure 4B) and verified extensive co-binding of these activators with SNAI1/2 distal peaks (Figure S3C). We next examined how TF binding changed upon SNAI1/2 depletion as a function of distance to the nearest SNAI1/2 motif (Figure 4B). We observed a strong distance-dependent effect: binding of all three TFs increased most sharply when their motif was within ∼200 bp of a SNAI1/2 motif, with the effect progressively diminishing beyond that distance (Figures 4C and 4D). This indicates that SNAI1/2 interfere with activators binding in their vicinity.

Interestingly, the mechanism may differ across TFs. The TWIST1 binding motif in CNCCs — a composite element called the "Coordinator" motif — contains both an E-box and a homeodomain motif.^41^ SNAI1/2 could therefore interfere with TWIST1 recruitment directly, by binding the shared E-box, as previously suggested in the case of the E-box binder MyoD^55^ (Figure S3D). NR2F1 and TFAP2A, however, bind motifs that are distinct in sequence from the SNAI1/2 E-box. The fact that their binding is nonetheless affected by SNAI1/2 depletion suggests that SNAI1/2 can act locally through a mechanism other than direct competition between TFs for a shared binding site.

Overall, these results suggest that the observed changes in H3K27ac and ATAC-seq signal reflect the combined action of multiple activators whose binding increases upon SNAI1/2 depletion, and which may raise acetylation levels by recruiting acetyltransferase cofactors. Indeed, a simple linear model predicting the change in H3K27ac upon SNAI1/2 depletion at enhancers bound by TWIST1, NR2F1 and TFAP2A shows that while TWIST1 binding change is the strongest contributor — consistent with the major role of TWIST1 in CNCCs^41^ — NR2F1 and TFAP2A each improve the model’s predictive power, suggesting that the H3K27ac readout reflects the combined effect of multiple TFs (Figure 4E).

### SNAI1/2 modulate heterogeneity of nucleosome occupancy at tuned enhancers

TFs constantly compete with nucleosomes for binding to DNA.^56^ The observation that SNAI1/2 depletion causes increased accessibility and activator binding, with delayed effects on H3K27ac, suggested a possible role for SNAI1/2 in limiting activator access to DNA through nucleosome remodeling. We therefore sought to use single-molecule chromatin profiling to examine nucleosome distributions at SNAI1/2-tuned enhancers and their changes upon SNAI1/2 depletion. We first applied Fiber-seq,^57^ in which nuclei are treated with an adenine methyltransferase (Hia5) that labels accessible, nucleosome-free DNA (Figure 5A). Long-read sequencing of the methylated DNA then reveals nucleosome footprints as ∼147 bp unmethylated patches within each individual molecule. Aggregating molecules into a metaplot of nucleosome midpoint density around distal, SNAI1/2-bound motifs, we found that nucleosomes were well positioned flanking the SNAI1/2 motif, with the dyad of the nearest nucleosome located ∼85 bp away (Figure 5B). SNAI1/2 depletion via dTAG reduced the signal of this proximal nucleosome, suggesting that SNAI1/2 contribute to its positioning.

**Figure 5:**
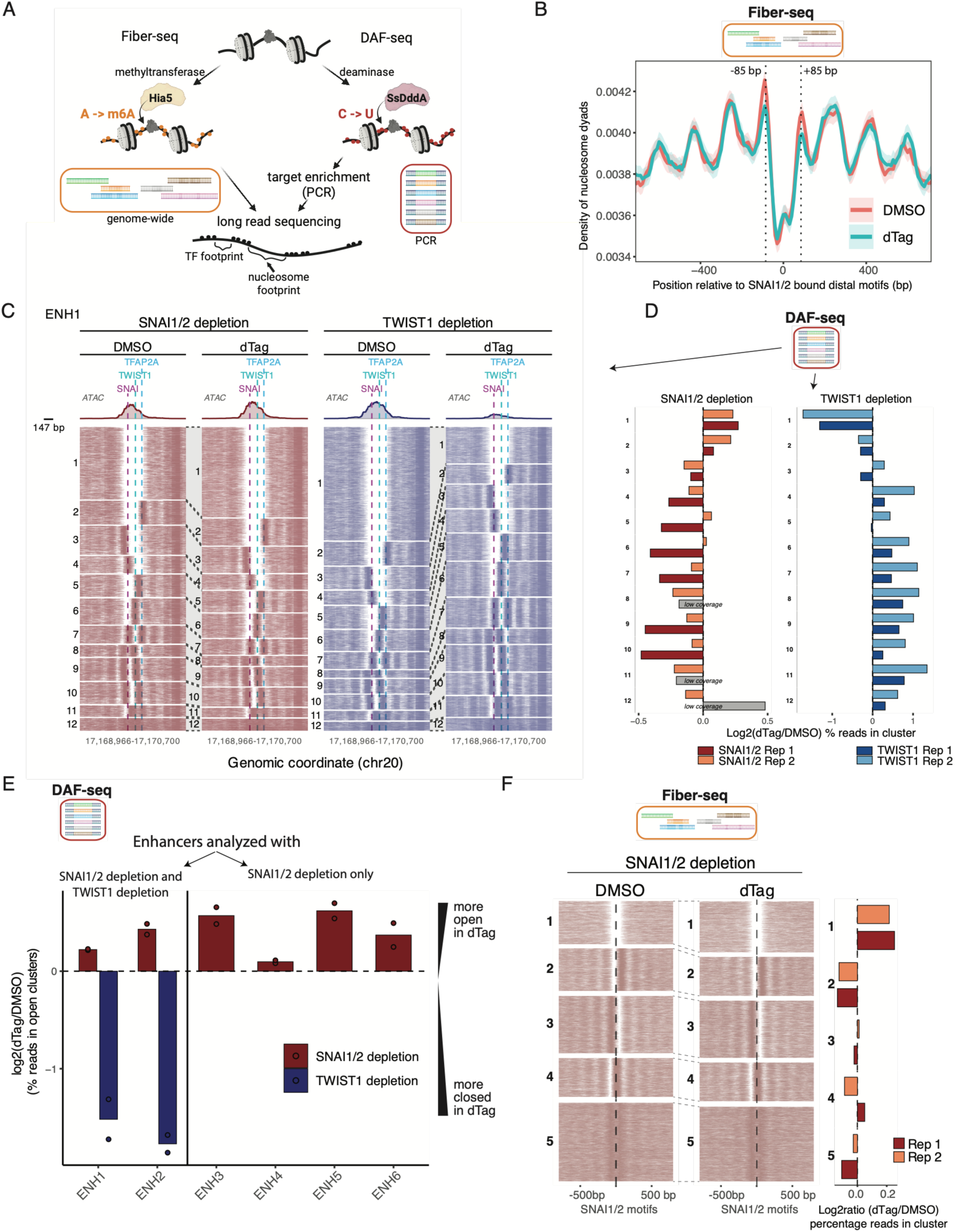
SNAI1/2 and TWIST1 shift the balance of heterogeneous enhancer chromatin states in opposite directions. (A) Schematic representation of Fiber-seq and DAF-seq protocol (created with BioRender.com). (B) Nucleosome midpoint density around distal SNAI2 bound motifs. Per-fiber nucleosome midpoint rate (nucleosomes per fiber per bp, mean ± SEM, 10-bp bins, 5-bin rolling mean) as a function of distance to the motif center, for DMSO- and 3 h dTAG-treated SNAI1-FNV SNAI2-KO CNCCs (two merged biological replicates each). Dotted lines, ±85 bp. (C) Single-fiber nucleosome heatmap of enhancer 1 showing DAF-seq data (two biological replicates per condition, SNAI1-FNV SNAI2-KO CNCCs and TWIST1-FV CNCCs, DMSO and 3 h dTAG for each line). Each row is one fiber; colored lines are nucleosome footprints (FiberHMM). Fibers were jointly k-means clustered (k = 12) on binary nucleosome occupancy within a region including 100 bp before the SNAI1/2 motif and 100 bp after the TWIST1 motif. ATAC-seq tracks are shown above the heatmap (DMSO and dTAG scaled together for each experiment). TWIST1-FV ATAC-seq data is publicly available^41^. 147 bp size scale is shown for visual reference. (D) Log2 ratio of the percentage of fibers assigned to each cluster in dTAG versus DMSO, shown separately for each biological replicate. Bars in grey represent clusters with <110 reads in both samples. (E) Bar plot showing summarized chromatin state redistribution for all six enhancers analyzed by DAF-seq. For each enhancer, fibers clustering by nucleosomes was performed as in C. Clusters were defined as “open” if having accessibility 150 bp before and after the SNAI1/2 motif (for red bars) or TWIST1 motif (for blue bars). Bars indicate replicates averages of the log2 ratio of the percentage of reads in “open” clusters in dTAG vs DMSO. Points are individual replicates. (F) Single-fiber nucleosome heatmap with Fiber-seq data (two biological replicates per condition). Shown are fibers overlapping SNAI2 bound motifs in tuned enhancers. Each row is one fiber; colored lines are nucleosome footprints (≥80 bp). Fibers were jointly k-means clustered (k = 5) on binary nucleosome occupancy within ±100 bp of the motif and ordered from least to most occupied. Right, log2 ratio of the percentage of fibers assigned to each cluster in dTAG versus DMSO, shown separately for each biological replicate.

Because Fiber-seq is performed genome-wide, it is difficult to achieve deep coverage at specific loci of interest. We therefore turned to Deaminase-Assisted Fiber-seq (DAF-seq),^58^ which uses a cytidine deaminase (SsDddA) to convert cytidines to uridines (Figure 5A). Because this conversion is encoded directly in the DNA sequence, it survives PCR amplification — allowing targeted, PCR-based enrichment of loci of interest without losing the underlying modification signal, in turn enabling deep analysis of thousands of molecules at individual loci. We amplified and sequenced one of the tuned enhancers previously validated by luciferase assay (Enhancer 1, Figures 3F-3H, Table S3). After calling nucleosome footprints with FiberHMM^59^ and clustering individual molecules based on nucleosome profiles around the accessibility peak, we found remarkable heterogeneity: fewer than half of the molecules contained an extended nucleosome-free region around the SNAI1/2 motif (we refer to these as “open” molecules or clusters for simplicity), while the rest displayed distinct nucleosome occupied patterns over the same interval (Figure 5C). The depth of our DAF-seq data allows clustering of distinct single-molecule chromatin fiber architectures, demonstrating that even within the defined mesenchymal CNCC cell state,^12^ individual enhancers occupy a heterogeneous landscape of architectures. Such heterogeneity of chromatin states at any given snapshot in time is suggestive of dynamic remodeling of enhancer chromatin.^60,61^

We next asked whether and how these patterns changed upon SNAI1/2 depletion. Since Enhancer 1 is also bound by the activator TWIST1 (Figure 4D), we used a previously generated FKBP12-V5-TWIST1 (hereafter TWIST1-FV) degron-tagged line^41^ to compare these changes with those occurring upon acute depletion of a strong activator (*i.e.,* TWIST1). Neither TWIST1 nor SNAI1/2 depletion led to the emergence of new chromatin states at this enhancer, as defined using DAF-seq-derived single-molecule chromatin fiber architectures. Rather, these perturbations shifted the rate at which individual architectures were used in opposite directions: acute TWIST1 depletion decreased the relative abundance of "open" molecules (*i.e.* with extended nucleosome-free region around the TF motifs), whereas SNAI1/2 depletion increased the abundance of open molecules and decreased that of nucleosome-occupied ones (Figures 5C and 5D). To extend this finding, we analyzed an additional enhancer with both SNAI1/2 and TWIST1 depletion (enhancer 2), plus four more enhancers with just SNAI1/2 depletion (enhancers 3,4,5,6). All examined cases recapitulated the heterogeneity of chromatin states observed at enhancer 1 and the effect of TF depletions: SNAI1/2 depletion increased the percentage of molecules with open conformations, while TWIST1 depletion decreased it (Figures 5E and S4, Table S3).

To further ensure that these observations were generalizable, we returned to the genome-wide Fiber-seq data and performed single-molecule clustering, centering molecules on SNAI1/2-bound motifs within tuned enhancers (defined as having increased H3K27ac upon 3 hours dTAG treatment). As observed with the DAF-seq data, SNAI1/2 depletion shifted the rate at which different chromatin states were occupied, increasing the abundance of open fibers (Figure 5F). Together, these results suggest that activator and repressor TFs act by tuning the frequency with which enhancers adopt each chromatin state, shifting the population-level distribution toward or away from open architectures.

### TF depletion modulates chromatin states of neighboring peaks within clustered enhancers

Mammalian enhancers are often organized in clusters of closely spaced nucleosome-depleted peaks. We reasoned that our DAF-seq data may provide insights into how loss of activators or repressors impacts chromatin fibers across peaks within such clustered arrangements. For example, enhancer 2 (ENH2) is composed of three distinct ATAC-seq peaks in close proximity to each other (368 or 313 base pairs summit to summit). Peak 1 is bound by TWIST1, peak 2 is bound by SNAI1/2 and TFAP2A, and peak 3 is bound by other unknown factor(s) (Figure 6A). This allowed us to investigate whether TWIST1’s and SNAI1/2’s effect on enhancer chromatin states is limited to the peak bound by these respective TFs or whether it can spread to neighboring elements within the clustered enhancer. We performed clustering of the single molecules based on each of the three ATAC-seq peaks separately and then compared changes in the fraction of open molecules at each peak upon depletion of either SNAI1/2 or TWIST1 (Figures 6B, 6C and S5A). As expected, both SNAI1/2 and TWIST1 depletion affected most strongly the region they directly bound (peak 2 or peak 1, respectively) (Figures 6B and 6C). However, a moderate effect in the same direction was also measurable at the neighboring peaks, indicating that TF depletion, while strongest in the immediate vicinity of the respective motif, can also influence chromatin state at distinct nearby elements.

**Figure 6:**
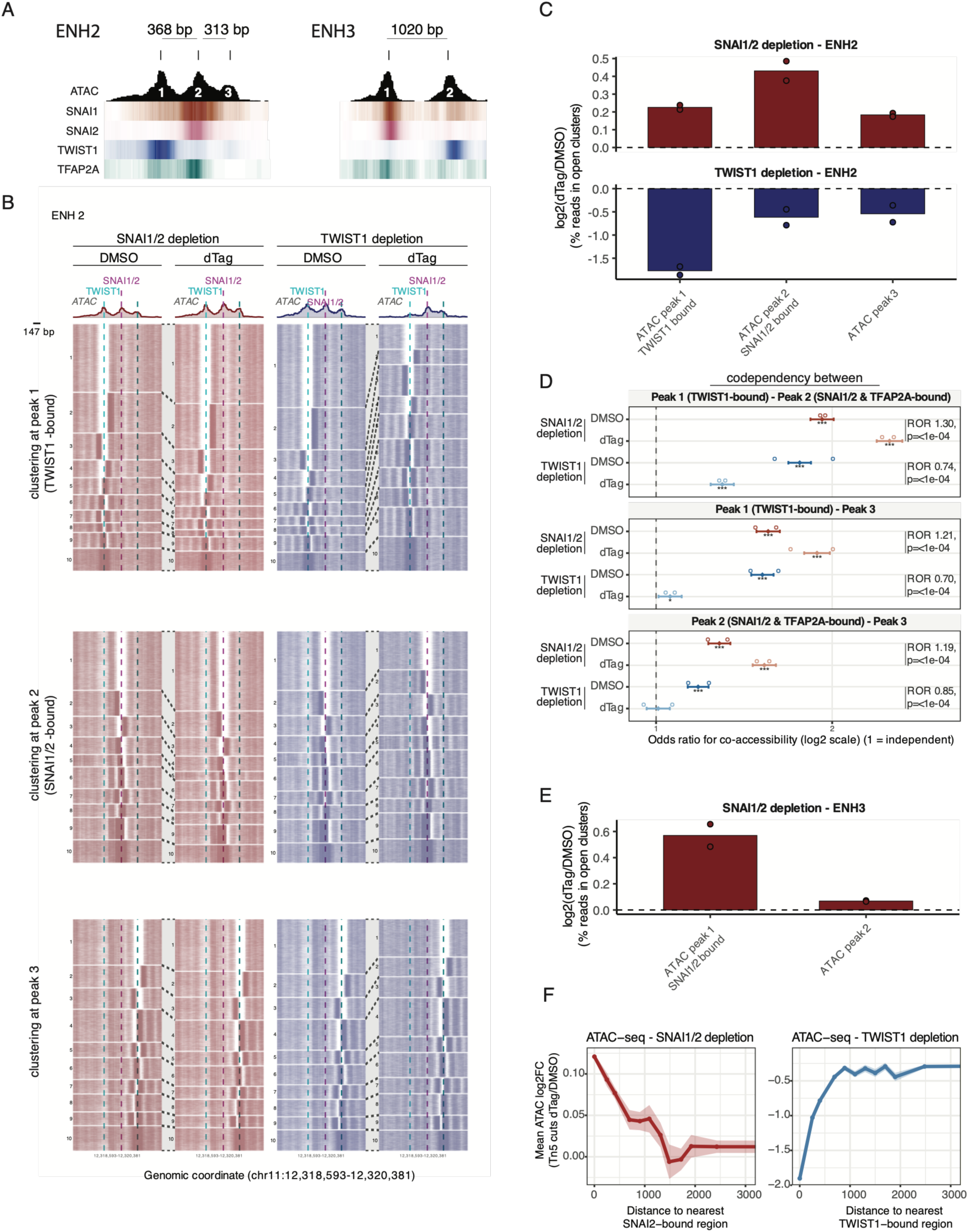
TF-depletion effects on chromatin state propagate to neighboring peaks within clustered enhancers. (A) ATAC-seq and ChIP-seq tracks showing the clustered arrangement of enhancers 2 and 3. (B) Single-fiber nucleosome heatmaps of enhancer 2 showing DAF-seq data (two biological replicates per condition, SNAI1-FNV SNAI2-KO CNCCs and TWIST1-FV CNCCs, DMSO and 3 h dTAG for each line). Fibers were jointly k-means clustered (k = 10) on binary nucleosome occupancy within a 200 bp region around the TWIST1 motif in peak 1 (top heatmaps), a 200 bp region around the SNAI1/2 motif in peak 2 (middle heatmaps), or a 200 bp region around an E-BOX motif (CAGCTG) in peak 3 (bottom heatmaps). ATAC-seq tracks are shown above the heatmap (DMSO and dTAG scaled together for each experiment). TWIST1-FV ATAC-seq data is publicly available^41^. (C) Bar plot showing summarized chromatin state redistribution for each neighboring peak at enhancers 2. Clusters (from panel B) were defined as “open” if having accessibility 150 bp before and after the motif used as reference for each peak. Bars indicate replicates averages of the log2 ratio of the percentage of reads in “open” clusters in dTAG vs DMSO. Red bars are for SNAI1-FNV SNAI2-KO data and blue bars for TWIST1-FV data. Points are individual replicates. (D) Odds ratio for co-accessibility between pairs of ENH2 summits on individual DAF-seq fibers. A fiber was scored accessible at a summit if an MSP ≥100 bp overlapped it; only fibers spanning all three summits were used. Filled diamonds, Mantel–Haenszel OR pooled across two biological replicates (stratified by replicate) with 95% CI; open circles, individual replicates (Fisher exact). Dashed line, OR = 1 (independence); OR > 1, summits open together more often than expected. x axis log2. Asterisks, BH-adjusted CMH p (* < 0.05, ** < 0.01, *** < 0.001). Brackets give the ratio of odds ratios (ROR = OR_dTAG/OR_DMSO) from a fiber-level logistic interaction model with replicate-specific intercepts, likelihood-ratio test, BH-adjusted. (E) Bar plot showing summarized chromatin state redistribution for each neighboring peak at enhancers 3. Clusters (from Figures S4 and S5C) were defined as “open” if having accessibility 150 bp before and after the motif used as reference for each peak. Bars indicate replicates averages of the log2 ratio of the percentage of reads in “open” clusters in dTAG vs DMSO. Points are individual replicates. (F) Log2 ratio (dTAG / DMSO) for ATAC-seq-derived Tn5 cut events counted at regions spanning 300 bp around each ATAC summit. Data is shown as binned averages ± SEM as a function of distance from the closest ATAC summit overlapping a SNAI1/2 (left) or TWIST1 (right) bound motif.

Consistent with this crosstalk, the three peaks showed positive codependency, measured by comparing the observed co-accessibility (fraction of molecules simultaneously accessible at both peaks) to the one expected under independence (product of each summit’s individual accessibility fraction) (Figures 6D and S5B, Table S4). The codependency between peak 1 and 3 appeared stronger than the one between peak 2 and 3, suggesting that this feature is not merely a function of distance between regions (Figures 6D and S5B). This agrees with a recent genome-wide study in *Drosophila*, which similarly found that codependency between regulatory elements is not explained by genomic distance alone.^62^ Interestingly, the effect size of the codependency (odds ratio) changed in opposite directions upon SNAI1/2 or TWIST1 depletion: loss of TWIST1 led to a drop in codependency, while loss of SNAI1/2 led to increased codependency, suggesting that these TFs can affect the communication between neighboring enhancer elements (Figures 6D and S5B).

We repeated the same analysis with enhancer 3 (ENH3), which has two peaks: one bound by SNAI1/2 and another one located 1020 bp away (Figure 6A). Here, both effects were weaker after SNAI1/2 depletion, possibly reflecting the larger distance between the peaks: the chromatin state change at the neighboring peak (measured as the change in the fraction of open reads) and the change in codependency (Figures 6E, S5C-S5E, Table S4). To investigate if the effects of TF depletions on the neighboring enhancers can be measured across the genome, we analyzed ATAC-seq data from acute SNAI1/2 depletion alongside ATAC-seq data from TWIST1 depletion.^41^ We plotted the fold change in accessibility (as number of Tn5 cut events) upon TF depletion at all ATAC-seq peak summits as a function of distance from the closest TF-bound peak. In both cases, the effect was almost entirely lost beyond ∼1.5 kb, consistent with our DAF-seq results at enhancers 2 and 3 (Figure 6F). Together, these results show that SNAI1/2 and TWIST1 act locally but can propagate their effect on chromatin opening to neighboring elements within roughly 1.5 kb. Thus, spacing between peaks within enhancer clusters may affect cluster behavior and responsiveness to changes in levels of individual activator and repressor TFs.

### SNAI1/2’s interference with activators is nucleosome-mediated

We reasoned that the single-molecule resolution experiments could allow us to formally test whether SNAI1/2 interferes with activator recruitment through direct competition for binding at neighboring motifs on the same DNA molecule, or alternatively, whether the competition instead arises from increased nucleosome deposition that occludes activator binding sites (Figure 7A). In the former scenario, SNAI1/2 depletion should increase activator binding even within nucleosome-free regions, whereas in the latter, it should only affect the fraction of nucleosome-free regions, but not the activator binding at already open sites.

**Figure 7:**
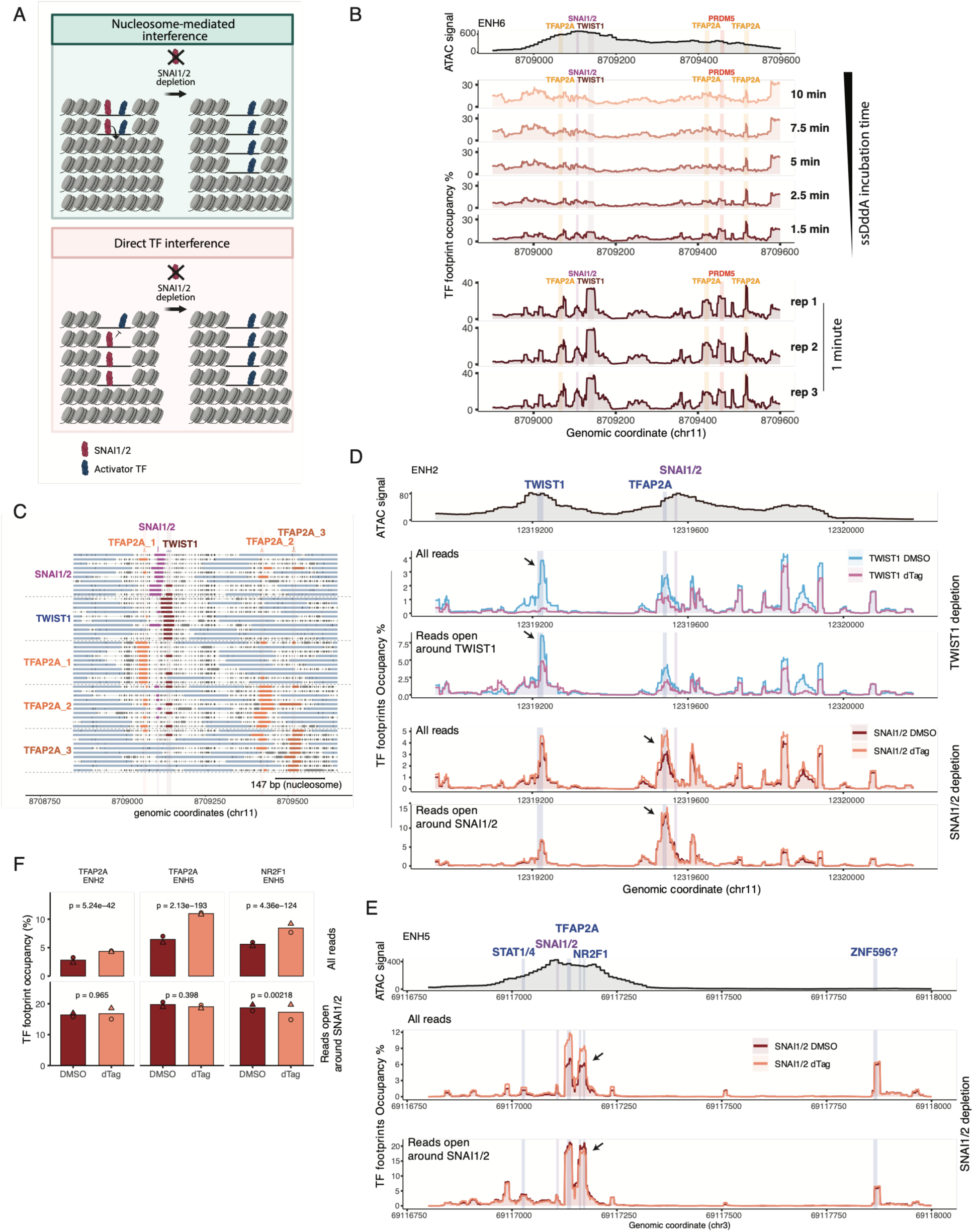
Single-molecule footprinting shows that SNAI1/2 interference with activators is nucleosome-mediated. (A) Visual representation of two possible models by which SNAI1/2 (red) could interfere with activators recruitment (blue). “Direct TF interference” changes the fraction of activator-bound molecules both on all molecules and only accessible molecules, while “nucleosome mediated interference” would not change the fraction of activator-bound molecules if looking at accessible molecules only. Schematic created with BioRender.com. (B) Metaplots of TF footprint occupancy percentage for enhancer 6, with the indicated SsDddA incubation times. An ATAC-seq track (DMSO) is shown above the metaplot for reference. (C) Visual representation of footprint calls and deamination at exemplary fibers. Each horizontal line is one read; grey ticks mark deaminated (accessible) bases, blue bars nucleosome footprints, and colored bars TF-scale footprints (≤40 bp), colored by the binding site they overlap (grey, unassigned). Reads are grouped by which TF footprint they carry (10 reads per group, labeled at left; dashed lines separate groups), and boxes above the reads mark the SNAI1/2, TWIST1 and TFAP2A motifs. Scale bar, 147 bp (one nucleosome). (D,E) Metaplots of TF footprint occupancy percentage for enhancers 2 and 5, comparing DMSO and 3h dTAG conditions for either SNAI1-FNV SNAI2-KO or TWIST1-FV CNCCs. For each line, shown is a metaplot normalized over all reads and a metaplot of only reads belonging to clusters with extended nucleosome-free regions around the indicated TF motif (ENH2: clusters 1 and 2 for TWIST1, cluster 1 for SNAI1/2 [Figure 6B], ENH5: cluster 1 [Figure S4]). An ATAC-seq track (DMSO) is shown above the metaplot for reference. (F) TF footprint occupancy at individual activator motifs upon SNAI1/2 depletion. DAF-seq data from SNAI1-FNV SNAI2-KO cells treated with DMSO or 3 hours dTAG. Occupancy is the per-base fraction of reads carrying a TF footprint averaged across the motif. Top row, all reads at the locus; bottom row, only reads in the clusters with extended nucleosome-free region around SNAI1/2 (cluster 1 in each case). Bars show the mean of two biological replicates, points show individual replicates (paired design: circles, rep 1; triangles, rep 2). p values, Cochran–Mantel–Haenszel test on per-molecule footprint calls (each read fully spanning the motif scored as footprinted or not), stratified by replicate.

Distinguishing between these two possibilities requires detection of both nucleosomes and activator footprints on the chromatin fibers. However, the very short residence time of many TFs on DNA makes their footprints much harder to detect, because during the standard 10 min labeling reaction at 25°C, TFs unbind and allow deamination (or methylation) to occur at the exposed site. We reasoned that shorter incubation times could improve TF detection, as previously shown.^63^ We therefore performed a time-course experiment on enhancer 6, comparing incubation times from the standard 10 minutes down to 30 seconds. Strikingly, even just 30 seconds of incubation time produced measurable deamination (median rate 0.16), indicating that SsDddA’s rapid catalytic activity allows for such short labeling pulses (Figure S6A). After running FiberHMM,^59^ we observed that TF footprints became progressively better resolved as incubation time decreased, with times under 1.5 minutes yielding clear footprints at the targeted binding elements while preserving nucleosome detection (Figures 7B, 7C, S6B and S6C). We validated this footprint signal by depleting TWIST1 in TWIST1-FV cells with a 3 hours dTAG treatment, which flattened the footprint at enhancer 2 (Figure 7D). In addition, we validated that these shorter times permitted the confident identification of reads with “open” and nucleosome-dense chromatin states (Figure S6D).

We performed DAF-seq on SNAI1/2 degron tagged cells, with both five-minute and one-minute deaminase incubations, clustered the reads by nucleosome pattern, and generated TF footprint metaplots using either all reads or only those containing an extended nucleosome-free region overlapping the SNAI1/2 motif (open reads). Notably, we could not confidently detect a footprint for SNAI1/2 itself, potentially reflecting either that the residence time of this factor along accessible templates may be rapid, or that its occupancy may be juxtaposed immediately adjacent to or overlapping that of a nucleosome, as suggested by the nucleosome dyad metaplots (Figure 5B). Upon SNAI1/2 acute depletion (3 hours dTAG) we detected overall occupancy increases at several nearby footprints overlapping known TF motifs including those for TWIST1, TFAP2A and NR2F1 (Figures 7D, 7E, 7F, S6E, and S6F). TF occupancy increased most strongly in the vicinity of SNAI1/2 motifs (Figures 7D, 7E, and S6F), mirroring our earlier observations of distance-dependent effects on TF binding measured by ChIP-seq (Figure 4C).

Strikingly, however, when we restricted the analysis to reads with extended nucleosome-free regions overlapping the SNAI1/2 motif, the effect of SNAI1/2 loss on occupancy of neighboring TFs disappeared (Figures 7D, 7E, 7F, S6E, and S6F). As a control, when we degraded TWIST1, its footprint diminished both across all reads and on open reads (Figure 7D). Together, these observations indicate that binding of activators to accessible DNA is not affected by the presence of SNAI1/2. Instead, our results are consistent with a model whereby SNAI1/2 tune enhancer activity through TF interference mediated by the increased frequency of the nucleosome-occupied enhancer states (top model in Figure 7A).

## Discussion

Precise control of gene expression levels is increasingly recognized as central to establishing cellular identity and function. Dosage sensitivity is widespread: over 3,600 human genes are estimated to be haploinsufficient and/or triplosensitive, underscoring the need for tight regulation of gene dosage at both transcriptional and post-transcriptional levels.^64^ This need is especially acute during embryonic development, when differentiation programs must be precisely timed in response to signaling cues. Here, we describe a previously unrecognized mechanism of transcriptional tuning in which the sequence-specific transcription factors SNAI1 and SNAI2 negatively modulate enhancer activity by reducing the frequency with which enhancers adopt an open chromatin state. In turn, increased nucleosome occupancy leads to reduction in activator TF binding, H3K27ac signal, and activation potential. SNAI1/2 are expressed across a wide range of cell types of diverse embryonic origin, and the phenotypes associated with their loss in mice span, and extend well beyond, neural crest-derived tissues.^11,14,65–68^ This suggests that the enhancer-tuning mechanism described here may operate broadly in many other cellular and developmental contexts. SNAI1/2 are also unlikely to be the only sequence-specific repressors capable of this mode of regulation: deep learning models have identified motifs for additional candidates — including HIC1/2, BCL11A, YY1/2, and NFY — predicted to negatively modulate accessibility at the active enhancers to which they bind,^11^ pointing to a potentially much broader role for sequence-specific repressors in enhancer tuning.

Enhancer accessibility is not a static feature fixed at the time of cell fate specification; rather, it must be actively maintained through the competition between transcription factors/cofactors and nucleosomes.^6^ Recent advances in single-molecule chromatin profiling have offered a glimpse into the heterogeneity of TF occupancy and nucleosomal configurations at cis-regulatory elements, showing that co-occupancy by multiple activator TFs increases the frequency of open chromatin states.^57,58,69–76^ Here we demonstrate that repressor TFs have the opposite effect: they facilitate transition to a nucleosome-occupied state. Importantly, this repressor-mediated enhancer tuning has functional consequences for activity, as measured in reporter assays. Thus, enhancer activity reflects not only cooperation among activators in their competition with nucleosomes and the action of chromatin remodelers, but also a dynamic tug-of-war between activators and repressors, played out through their opposing effects on nucleosome occupancy. How repressors promote nucleosome-occupied states at enhancers remains to be established, but likely mechanisms involve either direct deposition of histones via histone chaperones and remodelers, or more indirect action — for instance via histone deacetylation, which could stabilize nucleosomes and render them more refractory to eviction by activators.

Single-molecule chromatin profiling by long-read sequencing provides a powerful opportunity to investigate codependency among distinct regulatory elements. Previous studies have identified widespread codependency between elements separated by tens of kilobases, attributable at least in part to three-dimensional chromatin interactions.^57–59,62,77^ Consistent with these findings, siRNA-mediated depletion of the transcription factor CLAMP in Drosophila was shown to alter chromatin accessibility at distal regulatory elements that are co-accessible with a CLAMP-bound element.^62^ Our data extend this observation to mammals, by showing that depletion of SNAI1/2 or TWIST1 alters accessibility at neighboring elements that are not directly bound by these factors. Moreover, the sequencing depth afforded by DAF-seq enables us to quantify changes in codependency scores, revealing that SNAI1/2 and TWIST1 depletion shifts these scores in opposite directions, with the activator increasing, and repressor decreasing, codependency across peaks. However, effects on chromatin accessibility across peaks decay rapidly once elements are spaced more than ∼1 kb apart. This short-range effect suggests locally acting mechanisms, for example, nucleosomal organization or modification propagating across neighboring elements, and/or a shared local biochemical environment, such as a biomolecular condensate. Regardless, clusters of closely spaced elements — such as those seen at super-enhancers or locus control regions — are a common feature of mammalian cis-regulatory landscapes.^78,79^ Our results suggest that organization of clustered enhancer elements and repressor binding may affect their TF perturbation responses and ability to act cooperatively.

## Acknowledgments

We thank Thomas Tullius for help with Fiber-seq and DAF-seq data analysis, Aaron Straight and Jacob Price Schwartz for consultation on single-molecule chromatin profiling and nanopore sequencing, Shiran Bar, Marko Dunjic, Grace Bower, Peter Sicong Wang, Gabriel Lopez, for critical input on the manuscript, and Wysocka lab members for stimulating discussions. Genome sequencing for human/chimpanzee line was done with the Stanford Genomics Core Facility. This research was supported by Howard Hughes Medical Institute, the Nomis Foundation, NIH R35 GM131757 award and California Institute for Regenerative Medicine DISC0-17487 award for J.W. L.I. is a Howard Hughes Medical Institute Fellow of the Damon Runyon Cancer Research Foundation (DRG 2505-23). K.J.B. is supported by the Propel Postdoctoral Scholars Program, Stanford School of Medicine. A.B.S. holds a Career Award for Medical Scientists from the Burroughs Wellcome Fund and is a Pew Biomedical Scholar. This work was supported, in part, by NIH grants 1DP5OD029630 and 1U01HG013744 to A.B.S.

## Author contributions

L.I. and J.W. conceived the study and designed the research; L.I. performed most experiments and analyzed most of the data; K.B. performed ChromBPNet analyses; T.S., performed human/chimpanzee cells haplotype phasing; B.J.M. and S.C.B. contributed to DAF-seq experimental design and performed library preparation; D.G.M. performed protein purification for Fiber-seq and DAF-seq; J.M. contributed to DAF-seq data analysis; S.T. performed sample processing for ATAC-seq; C.N. contributed to cell line generation; A.B.S. supervised Fiber-seq and DAF-seq experiments and analysis; J.W. supervised all the project; L.I. and J.W. wrote the paper; K.B., T.S., B.J.M., D.G.M. contributed to manuscript writing; all authors discussed and reviewed the manuscript.

## Data and materials availability

All newly generated data will be deposited to Gene Expression Omnibus.

Codes are available in github (https://github.com/luciaichino/SNAI1-2_CNCC) and will be deposited in Zenodo upon paper acceptance.

Plasmids generated in this study will be deposited in Addgene upon publication. All other reagents are available upon request.

## Materials and Methods

### Cell culture

Human embryonic stem cells (hESCs) H9 (WiCell, WA09) were cultured in feeder-free conditions, in mTeSR1 medium (Stem Cell Technologies, 85850) on Matrigel Growth Factor Reduced (GFR) Basement Membrane Matrix (Corning, 356231) and passaged using ReLeSR (Stem Cell Technologies, 05872). For genome editing and clonal expansion, cells were switched to mTeSR Plus medium (Stem Cell Technologies, 100-0276) but moved back to mTeSR1 before differentiation to CNCC. hESCs cells were fed every day for mTeSR1 or every 2 days for mTeSR Plus, and passaged every 4-6 days.

HEK293FT cells (Invitrogen, R70007) were cultured in DMEM high glucose medium with sodium pyruvate and L-glutamine, supplemented with 10% v/v FBS and 1x GlutaMAX, non-essential amino acids, and antibiotic/antimycotic.

The HL130 human/chimpanzee tetraploid hybrid cell line (C3649 + H20961) was kindly gifted by the Fraser lab^40^ and grown with the same culture conditions and protocols as the hESCs cells.

Routine mycoplasma tests were performed.

### Plasmids and cloning

AAV donor templates were cloned into the pAAV-GFP (Addgene plasmid # 32395) backbone by digesting pAAV-GFP with SpeI-HF (NEB, R3133S) and XbaI (NEB, R0145S) and using NEBuilder HiFi DNA Assembly (NEB, E2621L) with PCR products of ∼1 kb homology arms and tags (see primers for homology arms selection). In the case of SNAI2, a PAM site mutation was created in the primer. Flexible linkers (glycine-serine or glycine-alanine) of 5-11 aa were added in between the degron and epitope tags and the TF. Both SNAI1 and SNAI2 were tagged at the C-terminus.

For the luciferase reporter assay, enhancers of interest were cloned in the pGL3 luciferase reporter using NEBuilder HiFi DNA Assembly (NEB, E2621L) with PCR-amplified genomic DNA and pGL3 backbone digested with XhoI (NEB, R0146) and NheI-HF (NEB, R3131).

### Genome editing

Genome editing was performed via nucleofection of hESCs cells with CRISPR ribonucleoprotein complexes, using Adeno-Associated Virus (AAV) as donor DNA for homology directed repair.

#### AAV preparation

AAV preparation was achieved by transfecting HEK293FT cells with 22 µg of pDGM6 helper plasmid (Addgene plasmid # 110660), 6 µg of pAAV donor template plasmid, and 120 µg of polyethylenimine (Sigma-Aldrich, 408719) diluted in Opti-MEM (Gibco, 31985070) in 1 ml total volume per 15-cm plate (2 plates were used per construct). Twenty-four hours post-transfection cells were switched to slow-growth media (2% v/v FBS instead of 10%). Three days post-transfection cells were harvested and pelleted. AAVs were purified from cell pellets using AAVpro Purification Kit Midi (All Serotypes) (Takara, 6675) following manufacturer’s instructions.

#### Nucleofection

hESCs cells were treated with 10 µM Y-27632 (Stem Cell Technologies, 72304) for at least 2 h prior to nucleofection and then harvested as single cells with Accutase (Innovative Cell Technologies, AT104-500). Annealed guide RNAs were generated by resuspending crRNA XT and tracrRNA (Integrated DNA Technologies, IDT) in IDT duplex buffer to 200 µM each, mixing them 1:1, heating to 95°C for 5 min and then cooling slowly at room temperature for 15 min. 800,000 cells were nucleofected in 100 µl of Ingenio Electroporation Solution (Mirus Bio MIR 50114) containing RNPs made by 15 min room temperature incubation of 1.7 µl (17 µg) Alt-R S.p. HiFi Cas9 nuclease V3 (IDT) with 3.3 µl of 100 µM annealed guide RNAs. In the case of SNAI2 knockout, 2 µl of 100 µM ssDNA homology-directed repair (HDR) template was also included to facilitate generation of a premature stop codon (Table S5). Nucleofection was performed using the P3 Primary Cell 4D-Nucleofector X Kit L (Lonza, V4XP-3024) and the CA-137 program. In the case of homology directed repair, 5-10 µl of AAV preparation were added to the media right after nucleofection. Media was changed after 4 hours, maintaining 10 µM Y-27632 for at least two more days. Cells were cultured until nearing confluency, and then dissociated with Accutase and plated at low densities (500 cells per well of a 6-well plate). Resulting colonies were picked into 48-well plates and expanded. The clones were genotyped using QuickExtract (Lucigen, QE09050) for DNA extraction and PCR with a primer outside the homology arms (Table S5). Edited colonies were confirmed by genomic DNA extraction using the Monarch Genomic DNA Purification Kit (NEB, T3010S) and Sanger sequencing of purified PCR bands. For both SNAI1-FNV and SNAI2-FNV only heterozygous edited clones were obtained. Therefore, another round of editing was performed after expansion of a heterozygous clone to achieve homozygous tagging.

Guide RNAs used:

SNAI1-FNV: GAGGGAGCCTCGAGGGTCAG
SNAI2-FNV: GTCTCTCCTGCACAAACATG (PAM site mutation AGG to AGT)
SNAI2 KO: CGTTGAAATGCTTCTTGACC, AGCGGTAGTCCACACAGTGA

### CNCC differentiation and passaging

CNCC differentiation was achieved with a previously established protocol^12,39,44,45^ adapted for reproducible generation of neuroectoderm (NE) spheres with AggreWell, as described^82^. 3×10⁵ hESCs cells were seeded into each well of an AggreWell 800 24-well plate (Stem Cell Technologies, 34815) in 2 mL of CNCC induction medium, yielding approximately 1000 cells per NE sphere (two wells were used for each differentiation batch). The induction medium consisted of DMEM-F12 and Neurobasal combined at a 1:1 ratio, supplemented with 0.5X Gem21 NeuroPlex (with vitamin A; Gemini, 400-160), 0.5X N2 NeuroPlex (Gemini, 400-163), 1X antibiotic–antimycotic, 0.5X Glutamax, 20 ng/mL bFGF (PeproTech, 100-18B), 20 ng/mL EGF (PeproTech, AF-100-15), and 5 μg/mL bovine insulin (Gemini Bio-Products, 700-112P). During sphere formation, 75% of the medium was exchanged daily by careful pipetting. NE spheres were harvested on day 4 and transferred to 6-well plates (one well of AggreWell 800 into two wells of a 6-well plate, which is 150 NE spheres per well), each containing 2 mL of fresh induction medium. By day 7, spheres had adhered to the plate surface and migratory CNCCs had begun to emerge outward from the central NE core. Medium was refreshed at this point (day 7), and every two days onward. On days 11–13, depending on the batch, the migrated CNCCs were dissociated by brief treatment with Accutase diluted 1:1 with PBS, filtered twice with cell strainers to remove the NE spheres, and re-plated onto fibronectin-coated 6-well plates (7.5 µg/mL human fibronectin (Millipore, FC010-10MG), 1 mL per well) at a density of 2×10⁶ cells/well in CNCC maintenance medium (we refer to these cells as P1 CNCC). The CNCC maintenance medium was composed identically to the induction medium described above, but with 1 mg/mL Bovine Serum Albumin (BSA, Gemini Bio-Products, 700-104P) in place of insulin. Two days later, P1 CNCCs were split with 1:1 Accutase/PBS and re-seeded at 1:3 dilution onto fibronectin-coated 6-well plates. The day after splitting to P2 CNCCs, the medium was changed to CNCC-BC medium (CNCC maintenance medium supplemented with 1 ng/mL BMP2 (PeproTech, 120-02) and 3 μM CHIR-99021 (Selleck Chemicals, S2924). From this point forward, all subsequent passages were done by splitting up to 1:6 in CNCC-BC medium, every 2–4 days. All experiments were performed with P4 or P5 CNCCs.

### dTAG treatment

dTAGV-1 (Tocris, 6914/5) was dissolved in DMSO to a 500 μM stock and diluted in media to a final concentration of 500 nM immediately before addition to cell culture plates. For acute depletion time courses, treatments were staggered so that all time points were harvested simultaneously; an equivalent volume of DMSO (0.1% v/v final) was added to a control sample at the time point corresponding to initiation of the longest treatment. For the 3- and 4-day treatments, dTAGV-1 exposure was started in P3 CNCCs, and cells were passaged into P4 in the continued presence of dTAGV-1 prior to sample collection.

### Human / chimpanzee hybrid cells genome phasing

DNA was extracted from HL130 human/chimpanzee tetraploid hybrid ESC cell line (C3649 + H20961)^40^ using MagAttract HMW DNA kit (Qiagen, 67563) following manufacturer’s instruction but with the addition of 3 hours incubation at 55°C with 100 µg of proteinase K before addition of RNAse A. High-fidelity (HiFi) whole-genome sequencing (WGS) was performed by the Stanford Genomics Core Facility. High-molecular-weight (HMW) genomic DNA libraries were constructed using the SMRTbell Prep Kit 3.0 (PacBio, Cat. #102-182-700) following the manufacturer’s instructions. High-fidelity (HiFi) long-read sequencing was performed on a PacBio Revio system using one Revio SMRT Cell (from the Revio SMRT Cell Tray; PacBio, Cat. #102-202-200).

### Western blot

Cells were washed with cold PBS, lysed by incubation for 10 min on ice in RIPA buffer (50 mM Tris pH 7.4, 250 mM NaCl, 1% Igepal CA-630, 0.5% sodium deoxycholate, 0.1% SDS) with 1x cOmplete EDTA-free protease inhibitor cocktail (Roche, 11873580001), and sonicated for 4 cycles of 30s ON/30s OFF on high power using the Bioruptor Plus (Diagenode). Insoluble material was removed by centrifugation at >16,000 × g for 10 min at 4°C. The supernatant was quantified by BCA protein assay (Thermo, 23225) and then denatured by addition of 1x NuPAGE LDS Sample Buffer (Invitrogen, NP0007) and 100 mM DTT and heating to 95°C for 7 min. Samples were normalized by BCA quantifications and then loaded in 4-20% Novex Tris-glycine gels (Invitrogen) or 4-12% NuPAGE Bis-Tris gels (Invitrogen) and run in Tris-glycine buffer (25 mM Tris, 192 mM glycine, 0.1% SDS) or MOPS buffer (Invitrogen, NP0001). Gels were transferred onto PVDF membranes (Thermo Scientific) for 1.5 h at 350 mA in Tris-glycine buffer with 20% methanol, blocked with 5% milk in PBS with 0.1% Tween-20 (PBST) for 30 min at room temperature, and then incubated with primary antibody overnight at 4°C followed by horseradish peroxidase (HRP)-conjugated secondary antibody incubation for 1 h at room temperature, with 4 washes of PBST after each antibody incubation. Antibodies used: SNAI1 (Cell Signaling Technology, 3879S; 1:500), SNAI2 (Cell Signaling Technology, 9585S; 1:500), HSP90 HRP conjugated (Cell Signaling Technology, 79641S, 1:1000), beta-actin (Abcam, ab49900, 1:25000), Goat anti-rabbit HRP conjugated (Proteintech, SA00001-2, 1:10,000). Chemiluminescence was performed with Amersham enhanced chemiluminescence (ECL) Prime reagent (Cytiva, RPN2232) and imaged with an Amersham ImageQuant 800 (Amersham)

### Luciferase assay

CNCCs were plated at 100-125K cells/well in a 48-well plate and transfected right after adhesion to the fibronectin (∼15 minutes). CNCCs were transfected with a 6:1 ratio for FuGENE6:DNA (Promega, E2691), using 0.25 ng pRL Renilla control plasmid, 5 ng modified pGL3 reporter plasmid, and 44.25 ng carrier plasmid (pUC19) in 20 µl of Opti-MEM per well of a 48-well plate. Cells were lysed 24 h after transfection and assayed with the Dual-Luciferase Reporter Assay System (Promega, E1960).

Experiments were performed with technical quadruplicates (individual wells of cells transfected) and at least two independent batches for each enhancer. In this manuscript we report 4 enhancers out of 18 tested; the four reported enhancers are representative of SNAI1/2 depletion effects for all tested enhancers that had measurable basal activity level in the assay. Data is shown as log-transformed; each point represents a replicate, horizontal bar denotes the group mean and error bars denote ± 1 SD. For each enhancer construct, DMSO- and dTAG-treated samples were compared using an unpaired two-sample Welch’s t-test on log(Firefly/Renilla) values. Analyses were performed in R using ggplot2 and ggpubr. No correction for multiple comparisons was applied across constructs.

### RNA-seq

Cells were lysed directly on plate with TRIzol (Invitrogen, 15596018) (1 ml per 1 well 6 well plate) and stored at -80°C until processing. RNA extraction was performed with Direct-zol™ RNA Purification Kit, Miniprep kit (Zymo Research, R2052) including the in-column DNaseI digestion. RNA samples were processed for sequencing by Novogene. RNA sequencing libraries were prepared using a eukaryotic mRNA-seq (poly-A selection) protocol and sequenced by Novogene on an Illumina NovaSeq X Plus platform, generating paired-end 150 bp reads to a depth of ∼6 Gb (∼20 million read pairs) per sample.

### ATAC-seq

We adapted the Omni-ATAC protocol^83^ for our samples, omitting the DNase I treatment step prior to cell harvest and substituting Ampure XP beads (Beckman Coulter, A63881) for DNA cleanup. Cells were harvested with Accutase, counted on a Countess II (Invitrogen), and a 50,000-cell aliquot was pelleted by centrifugation at 500 × g for 5 min at 4°C. Pellets were resuspended for 3 min in cold lysis buffer — resuspension buffer (RSB: 10 mM Tris-HCl pH 7.4, 10 mM NaCl, 3 mM MgCl2) supplemented with 0.1% Igepal CA-630, 0.1% Tween-20, and 0.01% digitonin — and lysis was stopped by adding excess RSB containing 0.1% Tween-20. Nuclei were pelleted again (500 × g, 10 min, 4°C) and resuspended in a 50 µl transposition mix containing 25 µl TD buffer, 2.5 µl TD enzyme (Illumina, 20034197), 16.5 µl PBS, 0.01% digitonin, 0.1% Tween-20, and water. Transposition proceeded for 30 min at 37°C, after which tagmented DNA was purified with the DNA Clean & Concentrator-5 kit (Zymo, D4013) and eluted in 21 µl of 10 mM Tris-HCl, pH 8. Libraries were pre-amplified for five cycles using NEBNext Ultra II Q5 Master Mix (NEB, M0544): 72°C for 5 min, 98°C for 30 s, then five cycles of 98°C for 10 s, 63°C for 30 s, and 72°C for 1 min. A 5 µl aliquot of each pre-amplified reaction was used in a qPCR side-reaction (identical cycling parameters, minus the initial 72°C step) to calculate how many additional PCR cycles each sample required. The remaining 45 µl was then taken through that optimal cycle number and purified with two sequential double-sided Ampure XP selections (0.5x/1.3x, then 0.5x/1.0x bead ratios for the first and second addition, respectively). Final libraries were quantified using the Qubit dsDNA High Sensitivity assay (Invitrogen, Q33231), checked for size distribution on Tapestation D5000 and pooled prior to sequencing.

### Chromatin Immunoprecipitation sequencing (ChIP-seq)

For each sample, cells from one confluent 10-cm plate were fixed with 1% methanol-free formaldehyde (Pierce, 28908) diluted in PBS for 10 min at room temperature; crosslinking was then stopped by adding 2.5 M glycine to a final concentration of 125 mM and incubating for another 10 min. Fixed cells were rinsed with PBS, detached by scraping, and pelleted by centrifugation at 1350 × g for 5 min at 4°C. After an additional PBS wash, pellets were flash-frozen and stored at -80°C until use.

On the day of the experiment, pellets were thawed on ice for 30 min and taken through three sequential lysis steps with rotation, 10 min at 4°C each: first in 5 ml of lysis buffer 1 (50 mM HEPES-KOH pH 7.5, 140 mM NaCl, 1 mM EDTA, 10% glycerol, 0.5% Igepal CA-630, 0.25% Triton X-100, 1x cOmplete EDTA-free protease inhibitor cocktail [PIC], 1 mM PMSF), then in 5 ml of lysis buffer 2 (10 mM Tris-HCl pH 8, 200 mM NaCl, 1 mM EDTA, 0.5 mM EGTA, 1x PIC, 1 mM PMSF), and finally in 300 µl of lysis buffer 3 (10 mM Tris-HCl pH 8, 100 mM NaCl, 1 mM EDTA, 0.5 mM EGTA, 0.1% sodium deoxycholate, 0.5% N-lauroylsarcosine, 1x PIC, 1 mM PMSF). Chromatin was sheared with a Bioruptor Plus (Diagenode) for 10-15 cycles of 30 s on/30 s off at high power, diluted further with lysis buffer 3, and cleared by centrifuging at maximum speed for 10 min at 4°C.

Triton X-100 was then added to a final 1%, and a small aliquot of the cleared lysate was set aside to verify chromatin yield and fragment size: it was diluted in elution buffer (1% w/v SDS, 100 mM NaHCO3), treated with 200 mM NaCl and RNase A (Thermo, EN0531) for 1 h at 65°C, followed by proteinase K (Thermo, EO0492) for 1 h at 65°C, and the DNA was purified with the QIAquick PCR Purification Kit (Qiagen, 28106). This DNA was quantified with the Qubit dsDNA High Sensitivity kit, allowing the bulk of the sheared chromatin to be normalized across samples before setting up immunoprecipitations. Antibodies used, each at the indicated amount per ChIP, were: SNAI1 (Cell Signaling Technology, 3879S; 10 µg), SNAI2 (Cell Signaling Technology, 9585S; 10 µg), H3K27ac (Active Motif, 39133; 5 µg), H3K27me3 (Active Motif, 61017; 5 µg), H3K4me1 (Active motif, 39299; 5 µg), H3K9me3 (Abcam, ab8898; 5 µg), TWIST1 (Abcam, ab50887; 10 µg), AP-2α (Novus Biologicals, NB100-74359; 10 µg), NR2F1 (Perseus Proteomics, PP-H8132-00; 10 µg), and V5 (Abcam, ab15828; 10 µg). Input samples were set aside.

Chromatin-antibody mixtures (at least 25 µg of chromatin per IP) were incubated overnight, then captured for 4-6 h with 100 µl of Protein G beads (Invitrogen, 10004D). Beads were washed five times with RIPA wash buffer (50 mM HEPES-KOH pH 7.5, 500 mM LiCl, 1 mM EDTA, 1% Igepal CA-630, 0.7% w/v sodium deoxycholate) and once with 50 mM Tris-HCl pH 8, 10 mM EDTA, 50 mM NaCl, before elution in elution buffer (1% w/v SDS, 100 mM NaHCO3) for 30 min at 65°C.

Reverse crosslinking of the eluate proceeded overnight, with RNase A (Thermo Fisher Scientific, EN0531) added to 0.2 mg/mL, followed by a 4-h proteinase K digestion (0.2 mg/mL final; Thermo Scientific, EO0492); DNA was then recovered with the QIAquick PCR Purification Kit (Qiagen, 28106). Input samples were processed alongside. Sequencing libraries were generated from up to 50 ng of input or ChIP DNA with the NEBNext Ultra II DNA Library Prep Kit (NEB, E7645S), using a qPCR-determined optimal cycle number for each sample, and were purified with a post-PCR double-sided Ampure XP bead selection (0.5x/0.9x).

### Protein purification

#### Expression and purification of Hia5

DNA adenine methyltransferase Hia5 was expressed from pETHia5 in *E. coli* T7 Express cells and purified as reported previously,^57^ with the following minor modifications. Protein expression was induced with IPTG for approximately 20 h at 16°C. Following cell lysis and clarification, Hia5 was purified by batch Ni-NTA affinity chromatography and dialyzed overnight. The dialyzed protein was further purified by FPLC using a HiTrap SP HP cation-exchange column and eluted using a NaCl gradient. Purified Hia5 was concentrated, quantified by Bradford assay, and stored at −80°C. Adenine methyltransferase activity of purified Hia5 was confirmed by a gel-based assay as reported previously.^57^

#### Expression and purification of SsDddA

*Simiaoa sunii* DddA (SsDddA) was co-expressed with its cognate inhibitor SsDddI in E. coli BL21(DE3) and purified on an FPLC as reported previously^58^, with the following modifications. After binding to a HisTrap Excel column, the DddA–DddI complex was denatured on-column with 6 M guanidine-HCl, and DddA was subsequently renatured by gradual exchange into renaturation buffer. Renatured DddA was eluted with imidazole, concentrated and buffer-exchanged into storage buffer, quantified by Bradford assay, assessed purity on SDS-PAGE, and stored at −80°C. Cytosine deamination activity of purified SsDddA was confirmed using a coupled enzymatic assay with a double-stranded FAM-TATATATAACATTTAAATATAT-3BHQ1 probe and uracil DNA glycosylase (UDG) in a qPCR instrument. The reaction was incubated at 30°C, where DddA-mediated cytosine deamination generated uracil, which UDG subsequently excised to generate an abasic site. The reaction temperature was then increased to 60°C to induce cleavage at the abasic site, and the resulting fluorescence increase was measured. SsDddA activity was further validated by performing DAF-seq reactions.

#### Expression and purification of SsDddI

*Simiaoa sunii* DddI (SsDddI) was expressed from pColDuet_nHis_SsDddI in E. coli BL21(DE3) cells. Cells were grown in LB containing kanamycin, and protein expression was induced with IPTG for 18 h at 25°C. Cells were harvested by centrifugation and lysed by sonication. Following clarification, the soluble fraction was loaded onto an FPLC HisTrap Excel column, and DddI was eluted using an imidazole gradient. Purified DddI was concentrated and buffer-exchanged into storage buffer, quantified by Bradford assay, assessed purity on SDS-PAGE, and stored at −80°C. The inhibitory activity of SsDddI was confirmed by adding SsDddI at various molar ratios relative to SsDddA in the activity assay described above. SsDddI inhibitory activity was further validated by performing DAF-seq reactions. A detailed protocol for expressing and purifying SsDddA, SsDddI, and DAF-seq is available at: https://fiberseq.github.io/The-Guide-for-DAF-seq/protocol/protocol.html

### Fiber-seq

Fiber-seq was performed as previously described with the following modifications^57^. CNCCs were collected with accutase:PBS 1:1 and counted. Two million cells per condition were taken for nuclei extraction. Cells were washed with PBS and lysed on ice in 200 µL of cold Nuclei Extraction Buffer (20 mM HEPES–KOH, pH 7.5, 10 mM KCl, 0.1% Triton X-100, 20% Glycerol, 0.5 mM Spermidine [added fresh]). Cells were spun at 600 x g for 3 min at 4°C and pellets were gently resuspended in 75 µL per reaction of cold Reaction Buffer A (15 mM Tris (pH 8.0), 15 mM NaCl, 60 mM KCl, 1 mM EDTA (pH 8.0), 0.5 mM EGTA (pH 8.0), 0.5 mM Spermidine [added fresh]). Nuclei integrity was verified with Trypan Blue staining. 1,000,000 nuclei were transferred to a PCR tube and the reaction was performed for 10 min at 25°C, in 60 µL of 1× Reaction Buffer containing 1.5 µL of 32 mM SAM and 1 µL of Hia5 (100 U/µL). The reaction was stopped with 6 µL 10% SDS and vortexing. Volumes were brought up to 100 µL and then DNA extraction was done with NEB Monarch Spin gDNA purification kit (T3010S) following manufacturer’s instructions.

Library preparation was performed with Native Barcoding Kit 24 V14 (SQK-NBD114.24, Oxford Nanopore Technologies), and samples were sequenced with Promethion flow cell (FLO-PRO114M, Oxford Nanopore Technologies).

### DAF-seq

DAF-seq experiments were performed as previously described^58^ with minor variations. CNCCs were collected with accutase:PBS 1:1 and counted. Two million cells per condition were taken for nuclei extraction. Cells were washed with ice-cold PBS and resuspended on ice in 60 µL Buffer A (15 mM Tris-HCl pH 8.0, 15 mM NaCl, 60 mM KCl, 1 mM EDTA pH 8.0, 0.5 mM EGTA pH 8.0, 0.5 mM spermidine). 60 µL of 2× Lysis Buffer were added, cells were mixed by gently tapping and incubated for 10 min on ice (2× Lysis Buffer: 40 mM Tris-HCl, 300 mM NaCl, 6 mM MgCl₂, 0.1% (w/v) digitonin). Cells were spun at 350 × g for 5 min at 4 °C, resuspended in buffer A, and counted with Trypan Blue staining to verify nuclei integrity. 250,000 nuclei were taken for each reaction in PCR tubes, bringing volume to 47 µL of buffer A, and adding 1 µL of UNG inhibitor (NEB M0281) per reaction. The deamination reaction was achieved by adding 2 µL of 100 µM SsDddA (final 4 µM), quickly pipette mixing while moving to a thermocycler set to 25 °C. The reaction was stopped after the indicated about of time (1 or 5 minutes for the main experiments) by addition of 1 µL DddI (1000 µM, 5-molar excess) and rapid pipette mixing. Nuclei were moved to ice and 50 µL of PBS were added to bring the volume to 100 µL before DNA extraction with NEB Monarch Spin gDNA purification kit (T3010S) following manufacturer’s instructions with elution in 35 µL of water.

PCR reactions were performed with repliQa HiFi ToughMix (QuantaBio 95200-100), using 70 to 100ng of DNA per reaction and 0.3 µM each of forward and reverse primers (Table S5), in 50 µl final volume. Reactions were incubated with 30 cycles of 98°C for 10 seconds, X °C (annealing temperatures in Table S5) for 5 seconds, and 68°C for 1 minute. PCR reactions were purified with NucleoSpin Gel and PCR Clean-Up (Takara Bio, 740609.250) and eluted in 15 ul of water. Different PCR reactions performed on the same template DNA were pooled equimolarly before library preparation.

The time course DAF-seq experiments were sequenced with Plasmidsaurus Premium PCR service (Oxford Nanopore, R10.4.1). All other DAF-seq experiments were processed as follows. Each pool of PCRs was end-repaired, A-tailed, and ligated to a universal Y-adapter following the SMRTbell prep kit 3.0 protocol (PacBio, PN 102-141-700), then purified with a 1.8× SMRTbell bead cleanup (PacBio, PN 103-294-600). The Y-adapter was annealed from two HPLC-purified oligonucleotides (oligo 1: 5′-AAGCAGTGGTATCAACGCAGAGAACGCAGAGTACT-3′; oligo 2: 5′-/5Phos/GTACTCTGCGTTAGATCGGAAGAGCGTCGTGTAG-3′; IDT) mixed at equimolar ratio to 200 µM, heated to 94 °C for 2 min, cooled to room temperature, and diluted to 20 µM in IDT Duplex Buffer. Adapter-ligated libraries were amplified with RepliQa HiFi ToughMix (Quantabio, PN 95200) using primers FWD 5′-CTACACGACGCTCTTCCGATCT-3′ and REV 5′-AAGCAGTGGTATCAACGCAGAG-3′ (98 °C, 30 s; 10 × [98 °C, 10 s; 58.8 °C, 30 s; 68 °C, 20 s]; 68 °C, 5 min), then bead-purified and quantified (Qubit 1× HS dsDNA assay, ThermoFisher Q33231). To enable multiplexed concatenation, the 16 pools were divided into four groups of four, and each pool within a group was assigned to one of four array positions (A–D) by a position-specific "hook-addition" PCR following the Kinnex 8-fold PCR kit protocol (PacBio, PN 103-072-000; 98 °C, 3 min; 9 × [98 °C, 20 s; 68 °C, 30 s; 72 °C, 4 min]; 72 °C, 5 min). Positions A–C used the manufacturer-supplied position-specific primer mixes. Position D required a custom primer mix in place of the kit-supplied HQ mix, as this position pairs a non-standard position-specific forward primer with a common reverse primer (D_FWD: 5′-ACCTCCTCCUCCAGAAUCTACACGACGCTCTTCCGATCT-3′; Q_REV: 5′-AUGCACACAGCUACUAAGCAGTGGTATCAACGCAGAG-3′; both HPLC-purified, IDT). The four position-tagged reactions within each group were pooled in equal volume, bead-purified, and quantified. Pooled, position-tagged DNA (5 µg per array) was ligated to a Kinnex barcoded adapter (bcM0001–bcM0004), circularized into 4-mer concatemer arrays, treated to degrade non-circularized products, and bead-purified, all following the Kinnex concatenation kit protocol (PacBio, PN 103-072-000). Concatemer libraries were annealed, bound, and cleaned using the standard PacBio ABC procedure and sequenced on a PacBio Revio.

## QUANTIFICATION AND STATISTICAL ANALYSIS

### RNA-seq data analysis

#### Transcript quantification

Paired-end reads were quantified directly from raw FASTQ files using Salmon (v1.10.1) in mapping-based mode against a Salmon index built from the GRCh38 cDNA transcriptome. Library type was auto-detected (-l A), and Salmon’s built-in sequence-specific and fragment-GC bias correction models were enabled (--seqBias, --gcBias).

#### Differential expression analysis

Transcript-level quantifications were imported into R with tximport v1.30.0 and summarized to gene-level counts (countsFromAbundance = "lengthScaledTPM") using a GRCh38 transcript-to-gene mapping file. Gene-level counts were analyzed with DESeq2 v1.42.0, using a design that modeled treatment condition (DMSO, 3h, 1day, 3days) together with a combined clone/replicate term. Genes with fewer than 5 reads in at least 3 samples were filtered out prior to model fitting, and DMSO was set as the reference condition. Wald test contrasts were then extracted between each treatment timepoint and the DMSO control. For visualization, log2 fold-change estimates were additionally shrunk using the apeglm v1.24.0 method (lfcShrink), which reduces the noise associated with fold-change estimates for low-count genes.

### ATAC-seq data analysis

ATAC-seq data processing and alignment. Raw reads were assessed with FastQC before and after trimming. Reads were adapter- and quality-trimmed with Trim Galore (Babraham Institute; phred33 encoding, Q25 3′ quality cutoff, stringency 3, minimum length 20 bp), removing the Nextera Read 1 transposase adapter sequence (5′-CTGTCTCTTATA-3′). Trimmed paired-end reads were aligned to the hg38 genome with Bowtie2 v2.5.4 ^84^ (22 bp seed length, 0 mismatches allowed in the seed, maximum insert size 500 bp). Alignments were filtered with samtools^85^ v1.22.1 to retain only reads mapping uniquely (MAPQ ≥ 2) and as proper pairs, and PCR duplicates were removed with Picard MarkDuplicates (REMOVE_DUPLICATES=true); these deduplicated, uniquely mapped BAM files were used for all downstream analyses.

#### Tracks and peak calling

Genome browser tracks were generated from the deduplicated BAM files using deepTools bamCoverage (10 bp bins, RPKM-normalized). Peaks were called independently for each sample from the deduplicated, uniquely mapped BAM files using MACS2^86^ callpeak (BAM format, effective genome size "hs", q < 0.01, --call-summits), yielding narrowPeak files with sub-peak summit calls.

#### Consensus peak set

To obtain a single set of regions for cross-condition quantification, a consensus peak set was defined from the four vehicle (DMSO)-treated samples (two independent clones, two replicates each). Peaks were intersected pairwise across all four DMSO replicates (bedtools intersect -u), retaining only peaks present in all four. For each resulting overlapping peak region, the constituent sub-peak with the highest MACS2 summit score (-log10 p-value) was selected, and a fixed 2 kb window (±1 kb) was defined around that sub-peak’s summit. This produced 120120 candidate regions; after removing regions on non-canonical chromosomes/scaffolds, 119728 consensus accessible regions remained and were used for all downstream quantification. Some windows retain minor overlap due to closely spaced summits, which were not merged further.

#### Quantification and differential accessibility

For each of the 16 ATAC-seq libraries (2 clones × 4 timepoints [DMSO, 3h, 1day, 3days] × 2 replicates), raw read counts over the 119,728 consensus regions were obtained from the deduplicated, uniquely mapped BAM files using bedtools^87^ (v2.31.1) coverage (-counts, -sorted). Differential accessibility was assessed with DESeq2^88^ v1.42.0. The count matrix was filtered to retain regions with ≥5 reads in at least 4 samples, and a model was fit with the design ∼ clone + replicate + condition to account for clone- and replicate-level variation while testing the effect of treatment. Pairwise contrasts were extracted between each treatment timepoint and the DMSO control, and regions were ranked by Benjamini-Hochberg-adjusted p-value.

#### ATAC-seq changes by distance to TF-bound summits

This analysis was performed by first selecting a consensus set of ATAC-seq summits from the four DMSO control libraries. For each library, the absolute summit position of every MACS2^86^ narrowPeak entry was extracted (peak start + summit offset), keeping multi-summit peaks as separate points. Summit positions from all four libraries were pooled and clustered with bedtools cluster using a maximum distance of 50 bp. Only clusters containing a summit from all four libraries were retained as reproducible, and each was assigned the mean position of its constituent summits. Final regions were defined as 300 bp windows (±150 bp) centered on these reproducible summits.

Filtered alignments were used to generate single-base Tn5 insertion profiles. Using `samtools`^85^, reads were retained only if they were properly paired (SAM flag 0×2) and had a mapping quality of at least 30; unmapped, secondary, supplementary and duplicate-flagged alignments (flags 0×4, 0×100, 0×800 and 0×400) were discarded, as were reads mapping to the mitochondrial genome. To enrich for nucleosome-free chromatin, only fragments with an absolute insert size below 120 bp were kept. Each surviving read was reduced to its 5′ terminus and shifted by +4 bp on the plus strand and −5 bp on the minus strand to account for the Tn5 dimer binding footprint, yielding one 1-bp interval per insertion event. Insertion sites were then counted within the 300-bp windows centered on reproducible peak summits (see above) using `bedtools intersect -c` ^87^, producing one count file per sample.

Differential accessibility was assessed with DESeq2^88^ v1.42.0 (design ∼ clone + replicate + condition), retaining only summits with ≥5 reads in at least 4 samples. log2 fold changes were computed for each dTAG timepoint versus DMSO. SNAI2 motif instances were identified genome-wide (hg38) by FIMO^80^ using the SNAI2 DBD PWM (Jolma et al., 2013)^46^ and restricted to those falling within ±100 bp of a SNAI2 peak summit, yielding a set of SNAI2-bound motifs. Analysis was limited to distal ATAC summits (regions previously classified as "enhancer" or "other_distal"). Each distal summit was labeled motif-positive if it overlapped a SNAI2-bound motif, and for every summit the genomic distance from its center to the nearest motif-positive summit center was computed (GenomicRanges::distanceToNearest). Summits were then grouped into distance bins (0, 1–300, 300–600, 600–800, 800–1000, 1000–1200, 1200–1400, 1400–1600, 1600–1800, 1800–2000, 2000–3000, >3000 bp) and the mean ATAC log2 fold change (3 h dTAG / DMSO) ± SEM was plotted against the median distance of each bin. The same analysis was done with published ATAC-seq data from TWIST1-FV CNCCs with 3h dTAG or DMSO treatment (GSE230315, SRR24257841 SRR24257840 SRR24257838 SRR24257837)^41^, using the same summit regions as for SNAI1/2, but using the coordinator motifs^12^ and this paper’s TWIST1 ChIP-seq data to label TWIST1 bound regions.

### ChIP-seq data analysis

#### Read processing and alignment

ChIP-seq reads were processed and aligned to the hg38 genome using the same pipeline described above for ATAC-seq (FastQC, Trim Galore trimming, Bowtie2 alignment on hg38, and removal of non-uniquely-mapped reads and PCR duplicates), with one difference: rather than trimming the Nextera transposase adapter, the Illumina universal adapter sequence (5′-AGATCGGAAGAGC-3′) was used.

#### Genome browser tracks

bigWig tracks were generated from the deduplicated, uniquely mapped BAM files using deepTools bamCoverage (10 bp bins, RPKM-normalized).

#### Peak calling

For SNAI1 and SNAI2 ChIP-seq samples, peaks were called with MACS2^86^ callpeak (BAM format, effective genome size "hs", q < 0.01, --call-summits) using the corresponding sample from dTAG-treated cells — in which the target protein had been acutely degraded — as the control/background input for MACS2. Resulting peaks were then filtered to remove regions overlapping the ENCODE hg38 blacklist (v2) using bedtools intersect (-v). Finally, to remove peaks arising from non-specific chromatin background, peaks that also overlapped peaks called from the corresponding input sample were removed, again using bedtools intersect (-v). While SNAI1 and SNAI2 peaks were largely overlapping, we used SNAI2 (endogenous antibody) peaks for further analyses (heatmaps and scatterplots) because SNAI1 antibody generated more non-specific peaks, and V5 antibody generated weaker ChIP signal and lower peak numbers. For TWIST1, TFAP2A, and NR2F1 ChIP samples, peaks were called with MACS2 as described above but without control sample and filtered for blacklist and input peaks overlap. Only peaks overlapping between the two DMSO replicates were retained.

#### SNAI1 and SNAI2 ChIP visualization

The Figure 2 heatmap was generated with deeptools v3.5.6 computeMatrix reference-point with --referencePoint center option, and plotHeatmap with –sortUsingSamples 5. The scatterplot of SNAI1 and SNAI2 anti-V5 ChIP enrichment was generated with R package ggplot2 after calculating coverage at peaks with deeptools^89^ multiBigwigSummary BED-file.

#### Motif enrichment

Motif enrichment for each peak set was performed with MEME suite v5.5.9 Analysis of Motif Enrichment (AME)^90^ using as database HOCOMOCO^81^ v14 core motifs plus the Coordinator motif^12^ manually added.

#### TF activators binding versus motif distances analysis

The TWIST1, TFAP2A, and NR2F1 peak sets were concatenated and raw read counts were obtained from the deduplicated, uniquely mapped BAM files using bedtools^87^ (v2.31.1) coverage (-counts, -sorted). Log2 fold changes of differential ChIP enrichment in DMSO versus dTAG treated samples were obtained with DESeq2^88^ v1.42.0. Peaks for each factor were filtered to the top 80% by MACS2 signal value and restricted to canonical chromosomes, and each peak summit (defined from the MACS2 narrowPeak summit offset) was assigned the log2 fold change of its overlapping ChIP peak. DNA-binding motifs for each source TF were identified from a FIMO scan of the corresponding motif model from the Jolma 2013 database^46^ except for Coordinator motif^12^ (TFAP2A [this set includes both TFAP2A and TFAP2C motifs]: TFAP2A_DBD, TFAP2A_DBD_2, TFAP2A_DBD_3, TFAP2C_DBD, TFAP2C_DBD_2, TFAP2C_DBD_3, TFAP2C_full, TFAP2C_full_2, TFAP2C_full_3; NR2F1: NR2F1_DBD, NR2F1_DBD_2, and NR2F1_full; SNAI2: SNAI2_DBD); for NR2F1 and TFAP2A, overlapping motif matches were first consolidated by retaining only the best-scoring (lowest p-value) match within each cluster of overlapping motifs. For each peak summit, the nearest source-TF motif within 100 bp was identified and paired with its associated log2 fold change, and the distance from this motif to the nearest SNAI2 motif was calculated in a strand-aware manner relative to the orientation of the SNAI2 motif. Peaks were further restricted to those overlapping annotated distal regulatory elements (see regions annotation paragraph). Log2 fold change values were then plotted as a function of distance to the nearest SNAI2 motif using a Gaussian-kernel-weighted local mean (bandwidth = 100 bp) with a weighted standard error ribbon, computed over a ±1000 bp window in 25 bp steps.

#### H3K27ac quantification and differential enrichment

Raw read counts for H3K27ac ChIP-seq samples over the 119,728 consensus regions previously annotated from ATAC-seq data were obtained from the deduplicated, uniquely mapped BAM files using bedtools^87^ (v2.31.1) coverage (-counts, -sorted). Differential H3K27ac enrichment was assessed with DESeq2^88^ (v1.42.0). The count matrix was filtered to retain regions with ≥5 reads in at least 4 samples, and a model was fit with the design ∼ clone + replicate + condition to account for clone- and replicate-level variation while testing the effect of treatment. Pairwise contrasts were extracted between each treatment timepoint and the DMSO control, and regions were ranked by Benjamini-Hochberg-adjusted p-value.

#### Linear modelling of H3K27ac changes from TF binding changes

Differential TF binding was quantified with DESeq2^88^ (v1.42.0) as described above, using each set of overlapping DMSO peaks for the respective TF, yielding a log2 fold change per peak. Consensus ATAC-seq regions (described above) and all TF peak sets were restricted to the canonical chromosomes (autosomes, X and Y), discarding unplaced and alternate scaffolds. Each ATAC region was then annotated with, for each TF separately, the unweighted mean log2 fold change of all TF peaks overlapping it by at least 1 bp; regions with no overlapping peak for a given TF were left unassigned. Analysis was restricted to distal regions with an assigned log2 fold change for all three TFs, giving n = 2648 regions. On this set we fitted ordinary least-squares models predicting the H3K27ac log2 fold change from every subset of the three TF predictors (three single-predictor, three two-predictor and one three-predictor model), together with an intercept-only null model, for eight models in total. Predictive performance was assessed by 10-fold cross-validation blocked by chromosome. Chromosomes were assigned to folds by greedy largest-first packing so that folds were approximately balanced in region count, and every region on a given chromosome was therefore held out together. For each model we report the distribution of R² across the ten held-out blocks (each computed against the mean of its own test block) and, as the summary statistic, the pooled out-of-fold R², obtained by concatenating the held-out predictions from all folds and comparing them to the global mean. Analyses used R version 4.3.1 (2023-06-16) with GenomicRanges^91^ [v1.54.1] and the tidyverse^92^ [v2.0.0]; the random seed was fixed at 42.

### Correlation between H3K27ac ChIP-seq and ATAC

To test whether the relationship between depletion-induced changes in chromatin accessibility (ATAC-seq) and H3K27ac changed over the course of SNAI1/2 depletion, we fit a linear mixed-effects model predicting H3K27ac log2 fold-change from ATAC-seq log2 fold-change, time point (3 h, 1 d, 3 d) as a categorical factor, and their interaction, with a random intercept for genomic region to account for repeated measurement of the same regions across time points (`lme4::lmer`, formula: `ac_l2fc ∼ atac_l2fc * timepoint + (1|region)`). The analysis was restricted to distal (enhancer or weak enhancer), SNAI1/2-bound regions that were called up-regulated in ATAC-seq or H3K27ac at at least one time point (*n* = 1,172 regions). Region-specific random slopes were not estimable with at most three observations per region and were therefore not included. Significance of the interaction was assessed by likelihood-ratio test against a reduced model with a single common slope (`ac_l2fc ∼ atac_l2fc + timepoint + (1|region)`), both models refit by maximum likelihood; the interaction was highly significant (χ²(2) = 464.2, *p* < 2 × 10⁻¹⁶), indicating that the slope of the ATAC–H3K27ac relationship differed across time points. Estimated marginal slopes within each time point and pairwise comparisons between time points were computed with `emmeans::emtrends`; given the sample size, inference used asymptotic (normal-theory) degrees of freedom, and contrast *p*-values were adjusted by Tukey’s method. Slopes were additionally tested against a null of 1, corresponding to equal magnitude of H3K27ac and accessibility change. Pearson and Spearman correlations between ATAC and H3K27ac log2 fold-changes were computed independently within each time point as descriptive statistics. Full model output, variance components, slope estimates and correlations are reported in Table S2. Analyses used R v4.3.1, lme4 v1.1.35, lmerTest v3.1.3 and emmeans v 1.10.0.

### Human / chimpanzee hybrid genome phasing and analysis

#### Tetraploid genome phasing

Pacbio sequencing reads were assembled into contigs with `hifiasm` and resulting contigs were mapped to hg38 reference genome with `minimap2`, followed by SNPs calling with `bcftools mpileup` routine. SNP were exported into BED file with `bcftools view` and masked reference genome created with `bedtools maskfasta`.

#### ChIP-seq processing

Five replicates were performed for each ChIP. The ChIP-seq and input samples from human/chimpanzee tetraploid hybrid cells were mapped as described for human ChIP samples, but using an N-masked hg38 reference genome (masked for all polymorphisms identified from the PacBio data). Peak calling was performed with MACS2^86^ callpeak (BAM format, effective genome size "hs", q < 0.01, --call-summits, no control sample), removing regions overlapping the ENCODE hg38 blacklist (v2) using bedtools intersect (-v). For downstream analyses we selected SNAI2 peaks present in at least 3 of 5 replicates.

To split reads by species of origin, we used SNPsplit^93^. Because SNPsplit compares only two haplotypes/strains at a time, human-versus-chimpanzee splitting was performed in two independent rounds, using the phased, haplotype-resolved genotypes of the human (H20961) and chimpanzee (C3649): round 1 compared H20961 haplotype 1 (human) against C3649 haplotype 1 (chimpanzee), and round 2 compared H20961 haplotype 2 against C3649 haplotype 2. For each round, the haploid VCFs of the two haplotypes being compared were merged with bcftools, restricted to sites at which the two haplotypes carried different genotypes, and recoded as homozygous diploid genotypes with a dummy FI (filter) field, as required by `SNPsplit_genome_preparation`. `SNPsplit_genome_preparation` was then run in dual-hybrid mode on each of the two resulting SNP files to generate two N-masked hg38 genomes (hap1-masked and hap2-masked); each masked genome was indexed with bowtie2-build.

ChIP-seq reads from each sample were mapped independently to both dual-masked genomes, and SNPsplit was run on each alignment (`--paired`) using the corresponding hap1 or hap2 SNP file, assigning reads to genome1 (human) or genome2 (chimpanzee) in each round. The human-assigned BAMs from the two rounds were merged with samtools, as were the chimpanzee-assigned BAMs, and each merged BAM was deduplicated (samtools fixmate/markdup -r) and converted to paired-end FASTQ (samtools fastq). The fastq files were mapped again to genomes masked with all human SNPs (human reads) or with all chimpanzee SNPs (chimpanzee reads). Genome browser tracks for each species were generated from the resulting species-sorted BAMs using deepTools bamCoverage (10 bp bins, RPKM-normalized). Raw read counts over the 119,728 consensus regions previously annotated from ATAC-seq data were obtained from the species-sorted BAM files using bedtools^87^ (v2.31.1) coverage (-counts, - sorted).

To map SNAI2 motifs on each haplotype-specific genome, SNPs were extracted from each haplotype’s VCF with `bcftools view -v snps` and incorporated into the hg38 reference with `bcftools consensus`, yielding a haplotype-specific FASTA. FIMO (MEME suite) was then run on each haplotype-specific FASTA with `--max-stored-scores 10000000`. The SNAI2_DBD motifs from Jolma 2013 database^46^ was used. Motifs were grouped by each ATAC-accessible regions (defined above) with `bedtools intersect -loj`.

To compare human-chimpanzee differences in SNAI2 motif strength with differences in ChIP-seq signal, FIMO SNAI2_DBD p-values at ATAC-accessible regions (defined above) were compared across the four haplotype genomes. Following Prescott et al.^12^, a full outer join of the four FIMO outputs was performed on ATAC region and motif coordinates, with motif instances missing from a given haplotype conservatively assigned a p-value of 0.005. Motif instances were retained only if the p-value was identical between hap1 and hap2 within each species, and a human-versus-chimpanzee p-value log-ratio was calculated for each retained motif and averaged per ATAC region. In parallel, species-sorted ChIP-seq read counts over the same ATAC regions were CPM-normalized using each sample’s species-specific library size, normalized to the matched input (log2[(ChIP CPM+1)/(input CPM+1)]) per species, and expressed as a human-minus-chimpanzee log2 fold-change, averaged across the five replicates of each mark (H3K27ac, SNAI2). Analysis was restricted to all distal-classified ATAC regions overlapping SNAI2 ChIP-peaks peaks (called from ≥3 of 5 replicates), and the per-region mean SNAI2 motif p-value log-ratio was correlated (Pearson) with the mean human-versus-chimpanzee log2 fold-change in H3K27ac and SNAI2 ChIP-seq signal.

### Regions and peaks classification – promoter versus distal

#### Genomic region annotation and chromatin-state classification

Each ATAC consensus region was annotated for distance to the nearest canonical, protein-coding transcription start site (TSS), using Ensembl version 111 gene models retrieved via AnnotationHub and ensembldb (v3.10.1, v2.26.1)^94,95^; regions within 1 kb of a TSS were classified as TSS-proximal. Next, we used histone marks to reclassify putative promoters contaminating the TSS-distal list. H3K27ac and H3K27me3 ChIP-seq signal, generated from this study’s own DMSO-treated samples (BAM files merged across replicates prior to bigWig track generation), and H3K4me1 and H3K4me3 ChIP-seq signal from a publicly available dataset (Prescott et al.^12^; SRA accessions SRR2096413, SRR2096414, SRR2096417, SRR2096419), processed and aligned using the same pipeline described above, were quantified over the 2 kb ATAC consensus regions (and over an expanded 6 kb window in the case of H3K27me3), using deepTools multiBigwigSummary (BED-file mode, raw output counts). For each region, log2 ratios of H3K27ac/H3K27me3 and H3K4me1/H3K4me3 were calculated (with a pseudocount of 10 added to each signal value). We reclassified the TSS-distal regions in these categories: "enhancer" if log2[H3K4me1/H3K4me3] > -0.2 and (log2[H3K27ac/H3K27me3] ≥ 1), "weak enhancers" if log2[H3K4me1/H3K4me3] > -0.2 and -1 < log2[H3K27ac/H3K27me3] < 1, "H3K27me3-enriched" if log2[H3K27ac/H3K27me3] ≤ -1, “promoter” if log2[H3K4me1/H3K4me3] ≤ -0.2. For SNAI1/2 peaks analyses, these four categories were further collapsed into two composite classes: "enhancer" and "weak enhancers" regions were combined into a "distal" class, and all remaining regions were combined into a "promoter" class.

#### SNAI2 ChIP-seq peak classification

SNAI1/2 ChIP-seq peaks were assigned to the composite promoter or distal class of the ATAC consensus region(s) they overlapped. For peaks overlapping more than one consensus region, the peak was assigned to the region whose center was closest to the peak’s MACS2 summit position.

### Evolutionary constraint of SNAI1/2 peaks and motif occurrences

We asked whether SNAI1/2-bound distal regulatory elements, and the SNAI2 motifs within them, are under purifying selection, using two independent measures: cross-species conservation (phastCons scores computed on the 470-mammal Zoonomia alignment in hg38^47–49^) and the probability of negative selection within humans (LINSIGHT^50^). LINSIGHT combines human polymorphism with cross-species divergence and is available only for hg19. Four element sets were analysed: (1) the 3,877 distal SNAI ChIP-seq peaks; (2) a subset of those peaks overlapping enhancers that gain H3K27ac after 3 hr or 1 day of dTAG treatment (N = 834), which we term ‘regulated peaks’; (3) the 4,493 SNAI2 motif occurrences inside the distal peaks, identified with FIMO (p < 1e-4^80,96^) using the SNAI2 position weight matrix of Jolma et al. (2013)^46^; and (4) the 982 of those motif occurrences inside the 834 regulated peaks. Motif occurrences and background regions overlapping ENCODE blacklist regions^97^ or assembly gaps were removed. For phastCons we report, for each set, the fraction of aligned bases with a score of at least 0.8, pooled over elements. For LINSIGHT we report the mean of the per-element mean score. Elements with fewer than half of their bases aligned were excluded from both statistics.

Because the conservation of a region depends strongly on its sequence composition and genomic context, each element set was compared with a background matched on the features most likely to confound it. For peaks, the background was drawn from all other accessible regions in the same wild-type CNCCs, excluding promoters (GENCODE transcription start sites +/- 500 bp^98^) and the peaks themselves. Peaks were split into width deciles, and within each decile one accessible region per peak was drawn by propensity-score stratification (nullranges::matchRanges^99^) on GC content, CpG density, repeat-masked fraction and log_10_ distance to the nearest TSS. This was repeated 200 times. For motif occurrences, the background was chosen as follows: for every FIMO motif hit, an unbound distal genomic copy of the identical 9-mer word, sampled without replacement, was chosen. This was repeated 100 times. This sampling procedure holds the motif sequence itself constant and asks only whether the bound copies are more conserved than unbound ones. Each element subset was tested against a background of its own size. For example, the regulated peaks were tested against the background regions paired to those 834 peaks within the same 200 draws. All sets were lifted to hg19 for LINSIGHT scoring with liftOver (-minMatch=0.95; Kent et al., 2002), keeping only elements that mapped back to their hg38 position within 5 bp and retained their width.

For every element set, the observed statistic was compared with the distribution of the same statistic across background draws by a one-sided z-test (observed value against the mean and SD of the draws). To ask whether the regulated peak or motif subsets are more constrained than the corresponding ‘all distal’ parental set, we also compared each subset with 1,000 samples of the remaining parent elements drawn to match the subset’s composition (width and GC content for peaks, 9-mer word for motifs), using the same z-test. phastCons and LINSIGHT scores were extracted with bigWigAverageOverBed^100^; interval operations used BEDTools^87^. Statistical analysis and plotting were done in R.

### ChromBPNet analysis

#### 1) ChromBPNet model training, validation, and interpretation

ChromBPNet is a deep learning model that predicts chromatin accessibility such as ATAC-seq data from DNA sequence alone^51^. Through a two-step (two-model) training approach, the sequence biases of the Tn5 enzyme are eliminated, leaving model predictions that reflect unbiased chromatin accessibility information. ChromBPNet was installed and run exactly as described in the repository documentation (https://github.com/kundajelab/chrombpnet), using the hg38 genome and a pre-trained human bias model available from the Kundaje lab’s Zenodo (https://zenodo.org/records/7443683)^51^. We then trained two single-task ChromBPNet models using the ATAC-seq data (Tn5 cut site counts rather than whole-fragment coverage) collected from DMSO and 3 hour dTAG-treated human CNCCs. All models were trained using the identical, default architecture and across the same set of 192,174 peak regions, sized at 2,114 bp in length, which were curated using the reproducible MACS2-called ATAC-seq peaks from two replicates of two clonal SNAI1-FNV-SNAI2-KO lines (clone 1 and clone 2). For training, we provided a set of 342,859 non-peak regions that we generated using the ‘chrombpnet prep nonpeaks’ command line tool. During training, model validation and testing came from peaks localized across chromosomes 8 and 20 (validation set) and 1, 3, and 6 (test set), while the rest of the peaks were used for training. While both the counts and profile performance of the models using this approach was robust (Fold_0, Figures S1C and S1D), we next confirmed that the performance was stable and not driven by a particular chromosome hold-out scheme by shuffling the training, validation, and test region sets and training five more models (5-fold cross-validation, Figures S1C-S1E). The following chromosomes were used for the validation and test sets, in that order: chr11 & chr17; chr2, chr5, chr9 (Fold_1); chr14 & chr21; chr4, chr7, chr12 (Fold_2); chr18 & chr22; chr10, chr13, chr16 (Fold_3); chr2 & chr7; chr15, chr19, chrX (Fold_4); chr4 & chr13; chr8, chr14, chr17 (Fold_5).

To extract the base-resolution importance (contribution) scores, which represent the importance of each base for the model’s prediction of bias-removed chromatin accessibility, we used the ‘chrombpnet contribs_bw’ command line tool (applying DeepLIFT/DeepSHAP^51,101,102^) for each chrombpnet_nobias.h5 model. Because we were most interested in uncovering the sequences that predict the total chromatin accessibility across the window, we interpreted only the counts head of each ChromBPNet model and therefore did not extract or interpret profile contribution scores. We next used a modified TF-MoDISco codebase^103^, TF-MoDISco-lite (https://github.com/jmschrei/tfmodisco-lite), with the ‘-n 500000’ flag to cluster individual instances of sufficient counts contribution (seqlets) and discover motifs predictive, either negatively or positively, of chromatin accessibility in human CNCCs. The standard ‘modisco motifs’ command line tool restricts seqlet significance calling to a 400 bp window centered on each input region, which could discard true motif instances outside of this window, particularly for those patterns that are not primarily centrally localized. To avoid this, we performed seqlet calling over the full ChromBPNet input window while leaving all other parameters unmodified and matching the CLI defaults. We used the TF-MoDISco-lite algorithm on all models, but used only the DMSO model for determining which motifs to move forward with in downstream analysis. To do this, we used the summarized seqlet counts and contribution weight matrix (CWM) for each motif pattern and manually identified motifs based on prior knowledge, existing literature, and motif databases such as JASPAR (https://jaspar.elixir.no/). We chose not to move forward with patterns that we 1) could not identify or 2) did not possess greater than 250 seqlets.

#### 2) Motif mapping and curation

To map the TF-MoDISco-lite motif patterns back to the hg38 genome, we used FI-NeMo (https://github.com/kundajelab/Fi-NeMo), installed and run with default parameters as described in the repository documentation. This allowed us to consider both the motif sequence and contribution score when identifying sequence hits across our CNCC ATAC-seq peak set. Importantly, we decided to use only our DMSO CrhomBPNet model as our source model for motif mapping, that way we could map motifs from an unperturbed state and later determine the effects of SNAI2 depletion across those same motif instances. We further curated our mapped motifs by removing any redundant or palindromic patterns or those sub-motifs that were embedded within larger motifs (*e.g.*, *SNAI2* within a *Coordinator*). Total counts contribution for each motif instance was then calculated by summing the contribution coverage across the entire length of the mapped motif without any resizing of the motif coordinates, as previously described^104^. All motifs are summarized in Figure S1F with the number of instances in the final curated set, their CWM where the height of the base reflects its predictive importance, and average counts contribution score derived from each ChromBPNet model.

#### 3) *In silico* motif perturbations and predictions

We leveraged our trained ChromBPNet models to predict the bias-removed effects of *SNAI2* motif mutation on human CNCC chromatin accessibility. The *SNAI2* motifs used for *in silico* mutagenesis were selected based on their net negative counts contribution scores, their overlap with SNAI2 ChIP-seq data, and their overlap with presence across enhancers that gained chromatin accessibility upon SNAI2 degradation. For each motif instance, a 2,114 bp genomic window centered on the motif was extracted from the hg38 reference genome. Because the SNAI2 motif is short (7 bp), the motif was scrambled by iterative random resampling. Candidate replacement sequences were drawn base-by-base rom the local flanking base composition (estimated from the rest of the 2,114 bp window) until a candidate was found in which every position differed from the wildtype sequence and which no longer contained the motif’s most negatively contributing component (i.e, GGT). Paired wildtype and scrambled sequences were then one-hot encoded and passed through the trained, bias-corrected ChromBPNet models. Base-resolution predicted accessibility profiles (Tn5 cut site coverage) were generated across wildtype and *SNAI2* mutant sequences using DMSO and 3h dTAG models. Next, per-base DeepSHAP counts-contribution scores were computed for both the wildtype and mutant sequences using all models. Contribution tracks were visualized as per-base sequence-letter tracks and as difference-from-wildtype (mutant minus wildtype) tracks, allowing the direct comparison of how the local contribution landscape around each *SNAI2* motif changes upon either motif mutation or SNAI2 degradation.

### Fiber-seq analysis

Raw nanopore signal data were basecalled using Dorado (v1.0.2, Oxford Nanopore Technologies, https://github.com/nanoporetech/dorado) with the super-accuracy (sup) basecalling model, simultaneously calling 5-methylcytosine/5-hydroxymethylcytosine (5mC_5hmC) and N6-methyladenine (6mA) base modifications, without read trimming (--no-trim). Barcoded libraries were demultiplexed with dorado demux using the Native Barcoding Kit 24 V14 chemistry (SQK-NBD114-24). Demultiplexed reads were aligned to the human reference genome (hg38) with dorado aligner, retaining base modification tags in the output BAM files.

Modification calls were thresholded with modkit (v0.5.0, https://github.com/nanoporetech/modkit) call-mods, applying a filter percentile of 0.1 to exclude the lowest-confidence modification calls prior to downstream analysis; filtered BAM files were sorted and indexed with samtools^85^. Single-molecule nucleosome positioning was inferred from 6mA methylation patterns using fibertools^105^ (ft add-nucleosomes), which classifies nucleosome-occupied and MTase-accessible (linker) regions along each read based on m6A footprint density. ft center was used to find nucleosomes around SNAI2 motifs (Jolma 2013^46^) located within 100 bp from a SNAI1/2 ChIP-seq summit annotated as distal (as described above).

#### Nucleosome metaplots around SNAI2 motifs

Two biological replicates were pooled within each condition (DMSO, 3 h dTAG), after verifying that they generated the same result. For every nucleosome call, the midpoint was computed as the mean of its centered start and end coordinates. Midpoints falling within ±1000 bp of the motif center were assigned to 10-bp bins. Counts were tabulated per fiber per bin. For each condition, the nucleosome midpoint rate in each bin was calculated as the mean per-fiber count divided by the bin width (nucleosomes per fiber per bp), with the standard error of the mean taken across fibers and likewise divided by the bin width. Mean and SEM traces were smoothed with a centered 5-bin (50-bp) rolling mean; bins lacking a full smoothing window were dropped. Traces are plotted over ±650 bp with shaded ribbons denoting mean ± SEM. Analysis was performed in R using data.table, dplyr/tidyr, zoo, and ggplot2.

#### Single fiber nucleosomes heatmap

To select motifs in tuned enhancers, distal SNAI2 peaks were intersected with regions of increased H3K27ac upon 3 h dTAG treatment. These peaks were ranked by their H3K27ac log2 fold change (dTAG/DMSO) and split into tertiles (dplyr::ntile), and the top tertile (Q3), corresponding to the enhancers with the strongest H3K27ac gain, was retained. For each Q3 peak, SNAI2 motif instances within 100bp from a SNAI2 ChIP-seq summits were selected. Motif centers were used as the centering coordinates for ft center (fibertools). For each of the four libraries (DMSO and 3 h dTAG, two biological replicates), a binary occupancy matrix was built from nucleosome calls (≥80 bp size) over ±100 bp around the motif center in 10-bp bins (20 bins), with an entry set to 1 if any nucleosome overlapped that bin on that fiber and 0 otherwise. Matrices from all four libraries were row-concatenated and clustered jointly by k-means (k = 5, nstart = 10, maximum 100 iterations, seed 42) so that cluster definitions are shared across conditions and replicates. Clusters were relabeled by the rank of their mean occupancy, from cluster 1 (lowest nucleosome occupancy at the motif) to cluster 5 (highest). Nucleosome footprints (≥80 bp) were plotted over ±800 bp around the motif center; fibers were indexed within each cluster and panels faceted by cluster with heights proportional to cluster size. Replicates were pooled within condition to give merged DMSO and merged dTAG heatmaps. To quantify redistribution between conditions, the percentage of fibers assigned to each cluster was computed separately for every library, and the log2 ratio of dTAG to DMSO percentages was calculated within each biological replicate and plotted as a bar panel aligned to the heatmap facets. Analysis used R with *data.table, Matrix, dplyr/tidyr/purrr, ggplot2, patchwork*, and *ggrastr*.

### DAF-seq analysis

For the samples sequenced with PacBio, raw PacBio Kinnex concatemer arrays (4 positions each) were segmented with ‘skera split’, and split into per-pool BAM files, preserving per molecule tags (‘rq’, ‘np’, ‘ec’, ‘zm’, barcode tags, ‘RG’). Samples were processed with the DAF-QC-SMK pipeline https://github.com/StergachisLab/DAF-QC-SMK which selects full length consensus reads to remove PCR duplicates. For downstream analyses we used consensus reads with at least 3 reads per group for all enhancers except for enhancer 1, for which we used all consensus reads because of lower coverage (minimum=1). Putatively deaminated positions in consensus reads were masked by replacing the observed base with the corresponding IUPAC ambiguity code (Y for C→T strands, R for G→A strands) using a custom Python script (pysam), which also recorded the inferred strand of origin in the st BAM tag. Base quality scores and MD tags were preserved, and secondary and supplementary alignments were excluded.

These “decorated reads” were used as input for FiberHMM v2.9.7 standard processing^59^, using fiberhmm-call (--enzyme ddda -c 8 --seq pacbio --io-threads 16 --region-parallel --skip-scaffolds) and fiberhmm-extract (--block-scores --keep-bed). Bed files of TF, nucleosome, or MSP regions were used for plotting with custom scripts.

Time course experiments sequenced with Oxford Nanopore Technologies were processed directly with DAF-QC-SMK standard pipeline and fiberHMM as described above, without calling consensus reads.

#### 1. Single-fiber clustering and the linked dense plot

Nucleosome footprint calls from fiberHMM v2.9.7 for both 1 minute and 5 minutes SsDddA treated samples were parsed as plain text into per-fiber aligned spans and per-footprint genomic intervals. Fibers overlapping the clustering window (200 bp around the reference TF in all cases, except for enhancer 1, where we used 364 bp around both SNAI1/2 and TWIST1) were retained together with all their footprints. Replicates of the same condition were pooled into a single pseudo-sample so that all conditions were clustered jointly on one shared set of clusters. After clustering, each fiber’s cluster label was mapped back to its replicate of origin, so that all fold-change statistics are computed per replicate while the clustering itself is condition-agnostic. Each fiber was converted to a binary vector over the clustering window (1 if a nucleosome footprint covers that bp, 0 otherwise). Fibers from all samples were pooled into a single matrix and partitioned by k-means [k = 12 or 10; nstart = 200, iter.max = 200, fixed seed = 42]. Clusters were then relabeled in ascending order of mean nucleosome occupancy, so cluster 1 is the most nucleosome-depleted ("most open") and cluster k the most nucleosome-occupied ("most closed"). All samples are drawn in a single shared coordinate space as side-by-side lanes (inter-lane gap = 15% of window width) rather than as independent panels. Within a lane, one row is one fiber and each rectangle is one nucleosome footprint (rasterized at 300 dpi). Cluster bands are stacked in ranked order and each band’s height is proportional to that cluster’s share of that sample’s fibers, so every lane occupies the same total height regardless of sequencing depth. TF motif positions are shaded as vertical bands, and a per-sample bulk ATAC-seq track (bigWig, imported per bp over the plotting window) is drawn above each lane. For each sample, the percentage of that sample’s fibers falling into each cluster was computed, and for each replicate pair the log2 ratio of these percentages (dTAG / DMSO) was taken per cluster. Using percentages rather than counts normalizes for differences in per-sample depth. Clusters in which both the numerator and denominator sample had fewer than a minimum number of fibers were flagged as low-coverage, greyed out and labeled rather than removed [min_reads = 110 for ENH1].

#### 2. TF footprint metaplots

TF footprint calls (*_tf.bed) from the same fibers were filtered on fiberHMM’s TF footprint quality byte (blockTq) [≥ 70] and on footprint width [≤ 40 bp], and restricted to the plotting window. Each TF footprint inherited the cluster assignment of its parent fiber from the nucleosome clustering; footprints on fibers that received no cluster assignment were discarded. Occupancy was computed per bp as occupancy(bp) = (TF footprints covering bp / fibers covering bp) × 100. Two normalizations were used: for per-cluster curves the denominator is the fiber coverage of that cluster only, whereas for pooled ("all clusters") curves the denominator is the sample’s total fiber coverage. Curves were generated both across all fibers and restricted to the most-open cluster(s) [cluster 1, or 1 and 2, see figure legends]. Replicates were averaged per bp after normalization. For the nanopore data (deamination time course) we did not apply the TF footprint quality filter.

#### 3. TF footprint barplots

TF footprint occupancy (quality score ≥ 70, width ≤ 40 bp) was computed per base as the number of TF footprints covering that base divided by the number of reads spanning it, expressed as a percentage; the reported value for each motif is the mean of this quantity across the motif window. For all-read occupancy the denominator was every read spanning the base; for accessible-read occupancy both numerator and denominator were restricted to reads in the accessible clusters (as defined above). For statistical testing, each read fully spanning the motif was scored as footprinted or not footprinted, and DMSO was compared with dTAG by a Cochran–Mantel–Haenszel test stratified by biological replicate. Analyses were performed in R with *GenomicRanges* and the *tidyverse*.

#### 4. Open / closed fiber ratios

Ranked clusters were collapsed into "open" and "closed" groups. For each sample, the percentage of its fibers in each group was computed using that sample’s total fiber count in the region as the denominator, and log_2_(dTAG / DMSO) was taken per group per replicate pair. Group membership was assigned data-driven from methylation-sensitive patch (MSP) calls in the DMSO samples only. For each cluster, all MSPs overlapping the specified TF motif window were collected and their mean total width, mean extent upstream of the motif, and mean extent downstream of the motif were computed. A cluster was called open if mean MSP width ≥ 300 bp, upstream extent ≥ 150 bp, downstream extent ≥ 150 bp, and it was supported by ≥ 20 overlapping MSPs; all remaining clusters were called closed, so no cluster is left unclassified and clusters with sparse MSP support default to closed rather than being trusted. Requiring accessibility to extend on both sides of the motif, rather than merely overlapping it, avoids calling a cluster open on the basis of a short patch that happens to touch the motif. Classification was run separately per each motif.

#### 5. Peak codependency (single-fiber co-accessibility)

##### Input data and accessibility calls

Per-molecule accessible patches were taken from the recalled methyltransferase-sensitive patch (MSP) calls produced by fiberHMM v2.9.7 (using consensus reads with at least 3 reads per group). BED intervals were converted from 0-based half-open to 1-based inclusive coordinates, and enhancer summit positions were specified in the same 1-based convention (summits were defined as explained in “ATAC-seq changes by distance to TF bound summits”, ENH2: chr11:12319212/12319580/12319893, ENH3: chr20:57258315/57259335). A fiber was scored as accessible at a summit if at least one MSP overlapped that single-base position. Unless stated otherwise, only MSPs of at least 100 bp were considered; because this threshold filters patches and never fibers, the set of molecules entering the analysis is independent of it.

##### Analysis population

For each enhancer, the analysis was restricted to fibers whose aligned span covered every summit of that enhancer. Defining this population once, before any accessibility is scored, means that the marginal accessibility at each summit and the joint co-accessibility of any summit pair are estimated on exactly the same molecules; observed and expected co-accessibility are therefore directly comparable, as are all summit pairs within an enhancer.

##### Per-replicate codependency

All quantities were computed separately for each biological replicate. For each pair of summits (i, j) the per-fiber accessibility calls were cross-tabulated into a 2 × 2 contingency table with counts n11 (accessible at both), n10 and n01 (accessible at one summit only) and n00 (accessible at neither), summing to n fibers. Marginal accessibility was defined as p(i) = (n11 + n10) / n and p(j) = (n11 + n01) / n, the observed co-accessibility as n11 / n, and the co-accessibility expected under independence as p(i) × p(j). Codependency was defined as the difference

codependency = P(accessible at i and j) − P(accessible at i) × P(accessible at j),

so that positive values indicate that the two summits are open on the same molecule more often than chance, zero indicates independence, and negative values indicate mutually exclusive opening.

Significance was assessed with a two-sided Fisher’s exact test on the same 2 × 2 table. Because the expected value is derived from that table’s own margins, the null hypothesis of independence is exactly the null hypothesis that the observed co-accessibility equals the expected value; the Fisher p-value is therefore the test of that difference, and the conditional odds ratio with its 95% confidence interval is the accompanying effect size. Treating the expected value as a fixed external constant would be incorrect, since it is estimated from the same molecules and carries its own sampling error. These p-values were adjusted by the Benjamini–Hochberg procedure within each experimental group, across all replicate × summit-pair combinations of that group. A percentile 95% confidence interval on the codependency difference was additionally obtained from 2,000 bootstrap resamples of fibers drawn with replacement, recomputing both marginals and the joint probability within each resample; these are nominal 95% intervals and are not adjusted for multiplicity.

##### Combining replicates

Replicates were combined by stratified pooling of the 2 × 2 tables rather than by pooling fibers. Pooling molecules across replicates that differ in marginal accessibility can attenuate or even reverse the apparent direction of association (Simpson’s paradox), whereas stratification cannot. A common odds ratio per experimental group and summit pair was estimated with the Mantel–Haenszel estimator stratified by replicate, and the corresponding p-value from the asymptotic Cochran–Mantel–Haenszel test. These p-values were adjusted by the Benjamini– Hochberg procedure across all experimental group × summit-pair combinations of the enhancer, and the adjusted values are those reported as the primary test of codependency. Confidence intervals were computed from the Robins–Breslow–Greenland variance of the log odds ratio.

Homogeneity of the odds ratio across replicates was assessed with Woolf’s test. Woolf p-values are reported unadjusted, as a diagnostic of whether pooling is warranted rather than as an inferential test. The test was significant (p < 0.05) for 5 of 14 group–summit-pair combinations, indicating replicate-to-replicate variation in the magnitude of the association. The direction of the association was nevertheless consistent between replicates for every combination except one, in which the pooled odds ratio was itself indistinguishable from 1. Because each replicate contributes on the order of 10⁴ fibers, Woolf’s test is sensitive to differences in effect size that are small relative to the effects reported here; pooled estimates are therefore shown alongside the individual replicate odds ratios in the figures, and the full per-replicate contingency tables are provided in Table S4.

Effect of transcription factor degradation on summit coupling. Whether the coupling between two summits differs between DMSO and dTAG treatment is a question about an interaction, that is, a ratio of odds ratios (ROR = OR[dTAG] / OR[DMSO]), where 1 indicates that the association is unchanged, values below 1 indicate weaker coupling after degradation and values above 1 indicate stronger coupling.

The primary test was a fiber-level logistic regression fitted within each cell line and summit pair, modelling accessibility at one summit as a function of accessibility at the other, a replicate factor, and an explicit product of accessibility and treatment:

accessibility(j) ∼ accessibility(i) + replicate + accessibility(i) × dTAG.

Including replicate as a factor gives every replicate its own intercept, so replicate-to-replicate differences in baseline accessibility cannot be absorbed into the treatment effect. The coefficient on the product term is the log ratio of odds ratios, and it was tested by a likelihood-ratio test (1 degree of freedom) against the model omitting that term. Because the log odds ratio is symmetric in the two summits, the result does not depend on which summit is treated as the response. As an independent cross-check, a Wald test was performed on the difference of the two pooled Mantel–Haenszel log odds ratios, using their Robins–Breslow–Greenland standard errors; the two estimates of the ratio of odds ratios were required to agree to within 0.15 on the log scale, and the analysis halts otherwise. Both the interaction likelihood-ratio p-values and the Wald cross-check p-values were adjusted by the Benjamini–Hochberg procedure across all cell line × summit-pair combinations of the enhancer.

Because the odds ratio is not invariant to the marginal probabilities under a threshold model, and because the treatments shift marginal accessibility substantially, the DMSO-versus-dTAG comparison was accompanied by a sensitivity analysis based on the tetrachoric correlation. The tetrachoric correlation is the correlation of the latent bivariate normal that reproduces an observed 2 × 2 table given its own margins, and is therefore invariant to where the thresholds fall; it was computed for each replicate by solving P(X > h, Y > k; ρ) = n11/n numerically, with h and k the normal quantiles of the observed marginals, and averaged across replicates so that fibers were never pooled. A change in association was considered robust only if the tetrachoric correlation moved in the same direction as the odds ratio, and this was true for all comparisons (Table S4).

##### Robustness and quality control

The entire analysis was repeated at MSP width thresholds of none, 100, 150, 200 and 250 bp to confirm that the direction and significance of the reported associations are not an artifact of the patch-width filter, with the correction applied independently within each threshold. The bootstrap was seeded so that reported intervals are exactly reproducible. The two enhancers were analysed as independent experiments; no p-value was adjusted across enhancers.

##### Software and code availability

All analysis was performed in R using *dplyr, tidyr, readr, purrr, stringr, tibble, ggplot2, ggpubr, scales* and *openxlsx*; Fisher’s exact test, the Cochran–Mantel– Haenszel test and the logistic regressions used the base *stats* package.

##### Figure descriptions

Triangle plots show, for each experimental group, the mean codependency (observed minus expected co-accessibility) across replicates for every pair of summits, on a colour scale shared by all panels of a figure. Forest plots show the replicate-stratified Mantel– Haenszel odds ratio and its 95% confidence interval for each group (filled symbols) with the individual replicate odds ratios overlaid (open symbols), one panel per summit pair, on a log2 axis; the dashed line at 1 marks independence, and the bracket gives the ratio of odds ratios between DMSO and dTAG with its Benjamini–Hochberg-adjusted interaction p-value. Per-replicate values, contingency tables and all test statistics are provided in Table S4.

## Supplementary Information

**Table S1 – related to Figure 1**. Statistics of species assignment for ChIP-seq reads from human/chimpanzee tetraploid hybrid cells.

**Table S2 – related to Figure 3E**. Linear mixed-effects model results predicting H3K27ac log₂ fold-change from ATAC-seq log₂ fold-change.

**Table S3 – related to Figures 5-7**. DAF-seq sequencing depth statistics per sample (consensus reads).

**Table S4 – related to Figure 6**. Statistics of codependency analysis from DAF-seq data.

**Table S5 – related to Materials and Methods.** All oligonucleotides used in this study.

**Supplementary Figure 1.**
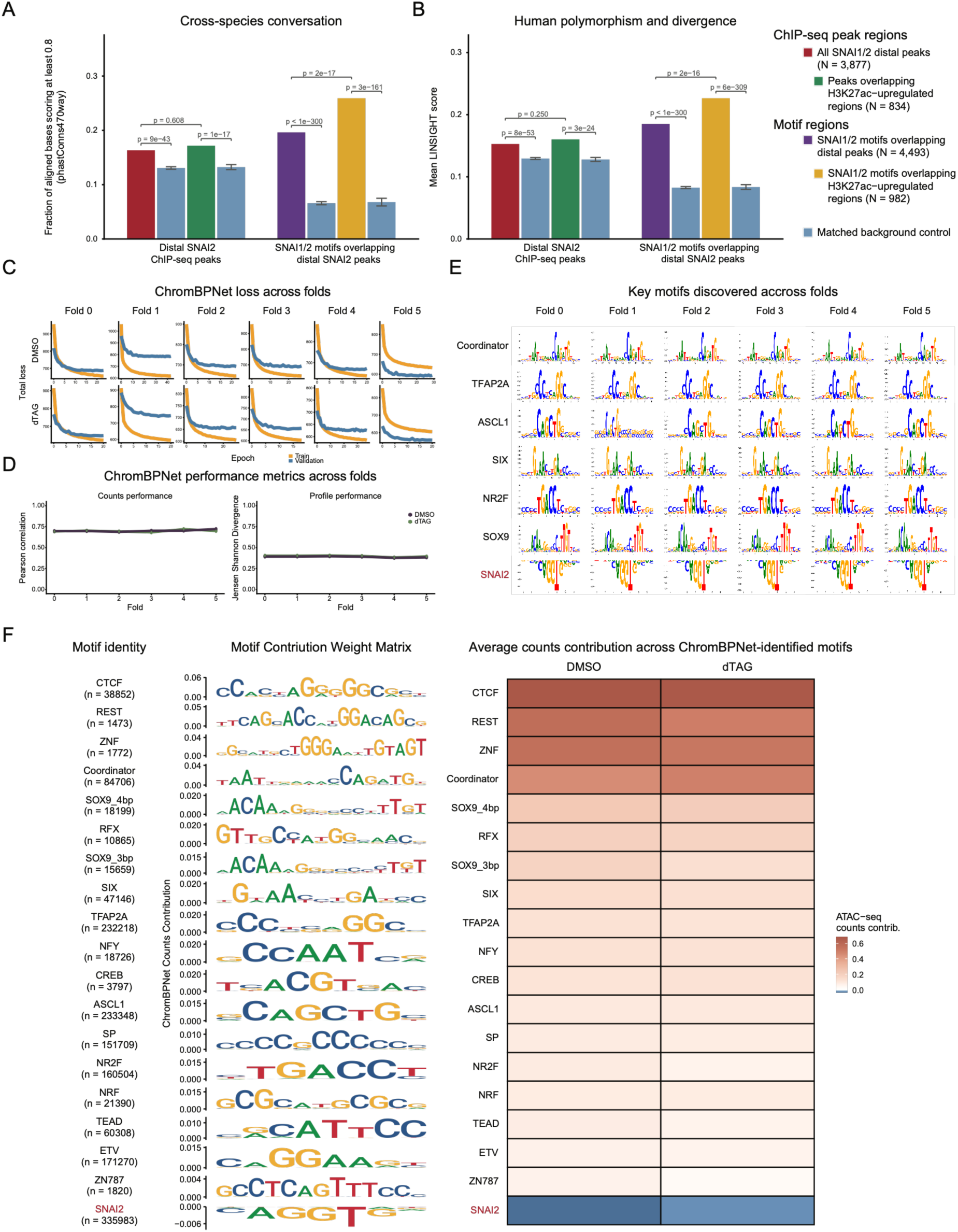
related to Figure 1: ChromBPNet model performance and learned motif syntax in human CNCCs. A) Fraction of aligned bases with phastCons470way >= 0.8, a measure of conservation across 470 mammals, for SNAI1/2 distal ChIP-seq peaks (red; N = 3,877), the regulated peaks, i.e. the subset overlapping enhancers that gain H3K27ac after 3 h or 1 day of dTAG treatment (green; N = 834), SNAI2 motif occurrences inside the peaks identified by FIMO at p < 1e-4 (purple; N = 4,493), and the motif occurrences inside the regulated peaks (yellow; N = 982). Blue bars show the background matched to the set immediately to their left, as mean and SD across background draws (see Materials and Methods). P-values above each pair are one-sided z-tests of the observed value against the mean and SD of its background draws. The P-value spanning the two foreground bars of each group compares the regulated subset (peaks or motifs at enhancers that gain H3K27ac) with 1,000 composition-matched samples of the remaining elements of its parent set (all distal peaks or motifs), and asks whether SNAI1/2 peaks in regulated enhancers are more constrained than all SNAI1/2 peaks. B) As in (A), the mean LINSIGHT score per element set and matched background. LINSIGHT estimates the probability that a point mutation at a site is deleterious by combining human polymorphism with cross-species divergence. C) ChromBPNet models were trained and validated across six data folds. Fold 0 is the interpreted model and folds 1-5 were trained for cross-validation purposes. The training (orange) and validation (blue) loss is consistent across all folds for both DMSO and 3 hour dTAG-treated (bottom) human CNCC ATAC-seq data. D) ChromBPNet counts and profile performance metrics are high across all data folds for DMSO (purple) and 3 hour dTAG (green) models. Performance metrics were evaluated across the same six folds as above and remained stable (∼0.7 for Pearson R and ∼0.4 for profile JSD), indicating that model performance is robust to the specific set of regions used for training, validation, and testing. E) The same key human CNCC motifs are discovered across all ChromBPNet data folds. Motif contribution weight matrix logos were generated by ChromBPNet’s courtesy TF-MoDISco-based overall_report.pdf output for each of the same six folds from above. While the number of associated seqlets differed across folds, key patterns were discovered irrespective of the regions into training, validation, and testing sets. F) ChromBPNet learns diverse sequence patterns predictive of chromatin accessibility in human CNCCs. For each motif, the FINEMO-mapped motif instances were curated and the corresponding contribution weight matrix was plotted to demonstrate the importance of each base within the motif. Across all mapped and curated instances, the average counts contribution score was determined and plotted from each of the single-task ChromBPNet models trained on DMSO and 3 hour dTAG-treated ATAC-seq data.

**Supplementary Figure 2.**
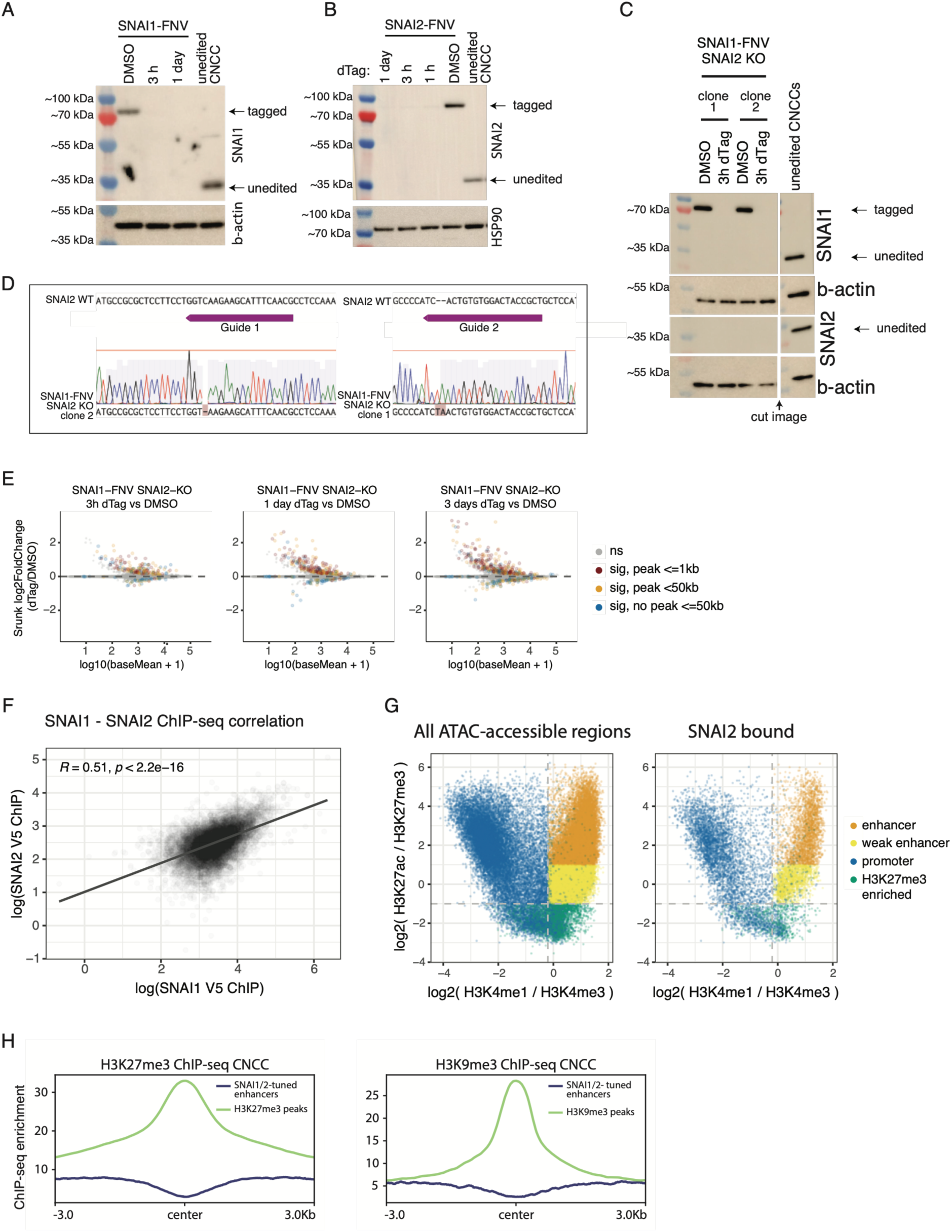
related to Figures 2 and 3: Generation and validation of SNAI1/SNAI2 degron lines and annotation of SNAI1/2 peaks. (A,B,C) Confirmation of SNAI1/2 tagging and depletion upon dTAG addition by western blot. (D) Sanger sequencing confirmation of SNAI2 KO (frameshift) mutations in two different clones. (E) MA plots of RNA-seq data for SNAI1-FNV SNAI2 KO lines for the indicated dTAG treatment time points. Red: significant genes with a SNAI1/2 ChIP-seq peak within 1kb, and no distal peak. Yellow: significant genes with a SNAI1/2 ChIP-seq peak within 50kb. Blue: significant genes with no SNAI1/2 ChIP-seq peaks within 50kb. Grey: not significant genes. (F) Scatterplot showing correlation between SNAI1 and SNAI2 anti V5 ChIP-seq at all SNAI1/2 peaks used in Figure 2C. Pearson R and p-value are shown. (G) Scatterplots showing classification based on histone marks for all 2kb regulatory elements used in subsequent analyses (left) or only the ones overlapping a SNAI1/2 peak (right). Enhancers were defined as not having a TSS within 1kb and also having log2(H3K4me/H3K4me3) > -0.2 and log2(H3K27ac/H3K27me3) > -1. H3K4me1 and H3K4me3 data are from Prescott et al.^12^ (H) Metaplots of H3K27me3 and H3K9me3 CNCC ChIP-seq enrichment at distal SNAI1/2-bound enhancers displaying increased H3K27ac upon 3h dTAG treatment, or at H3K27me3 and H3K9me3 ChIP-seq peaks as control.

**Supplementary Figure 3.**
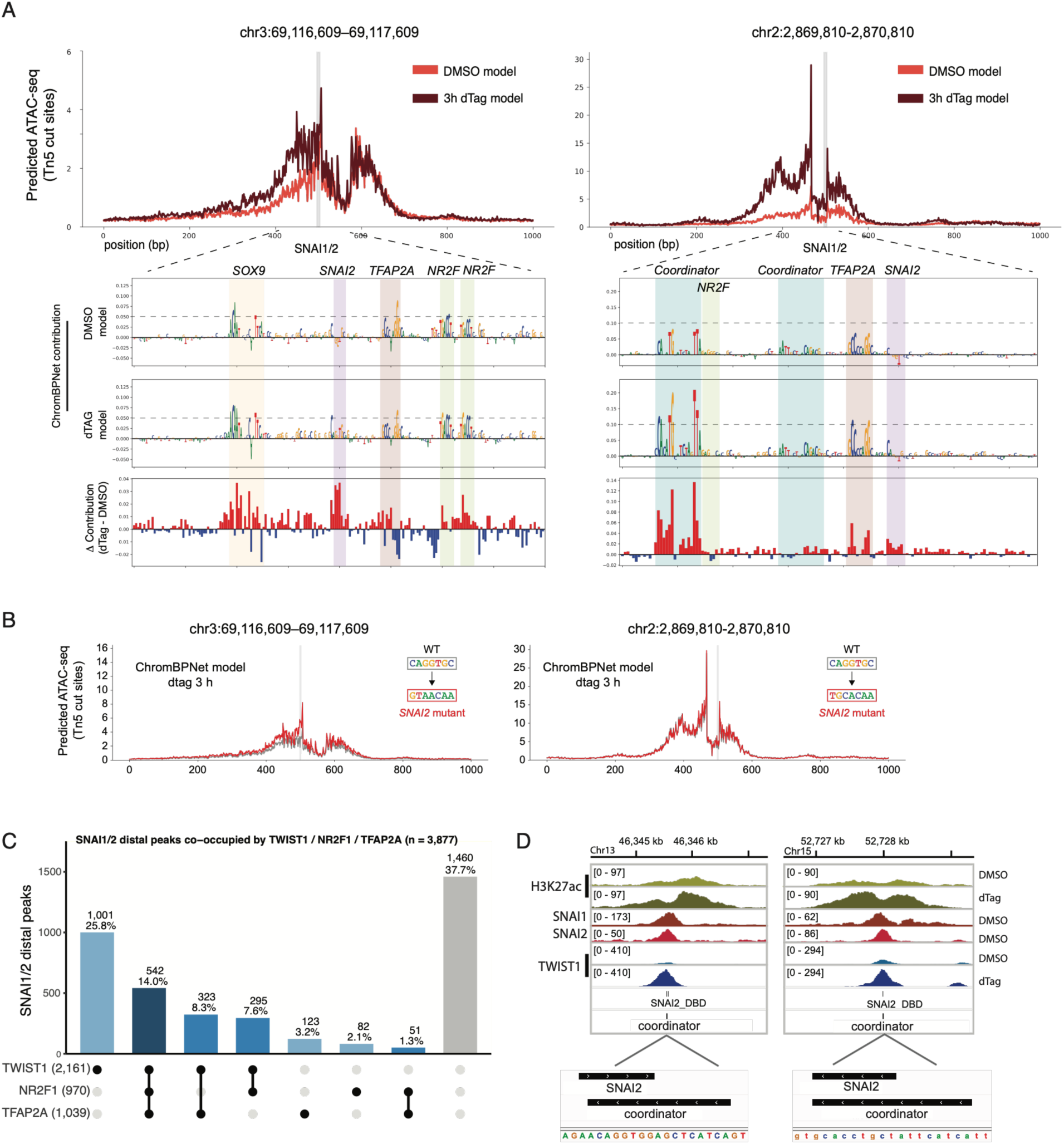
related to Figure 4: ChromBPNet captures loss of SNAI1/2 repressive contribution upon depletion and increased contribution at activator motifs. (A) ChromBPNet predicts increased accessibility across SNAI2-bound enhancers upon degradation of SNAI2 with a 3 hour treatment of dTAG. The DMSO and 3 hour dTAG-treated ChromBPNet models were used to predict chromatin accessibility across enhancers containing SNAI2 motifs. Upon degradation of SNAI2, ChromBPNet no longer attributes negative contribution to the SNAI2 motif, and activator motifs gain predictive importance (per base contribution tracks). (B) Tn5 cut site coverage predicted by the 3 hours dTAG depletion ChromBPNet model, across wildtype sequences (grey) and sequences in which the SNAI2 motif was mutated (red). (C) UpSet plot of SNAI1/2 distal peaks classified by the presence of TWIST1, NR2F1 and/or TFAP2A peaks within 500 bp (center-to-center). Filled dots and connecting lines below each bar indicate the factor combination. Bar labels give peak number and percentage of all SNAI1/2 distal peaks; shading reflects the number of co-occurring factors. Numbers in parentheses on the left are the total SNAI1/2 distal peaks within 500 bp of each factor (non-exclusive). (D) Examples of putative direct competition for binding to the same DNA region by TWIST1 and SNAI1/2. Shown are ChIP-seq normalized tracks for SNAI1-FNV SNAI2 KO (DMSO or 3h dTAG treatment). SNAI1/2 and TWIST1 overlapping motif annotations are highlighted.

**Supplementary Figure 4.**
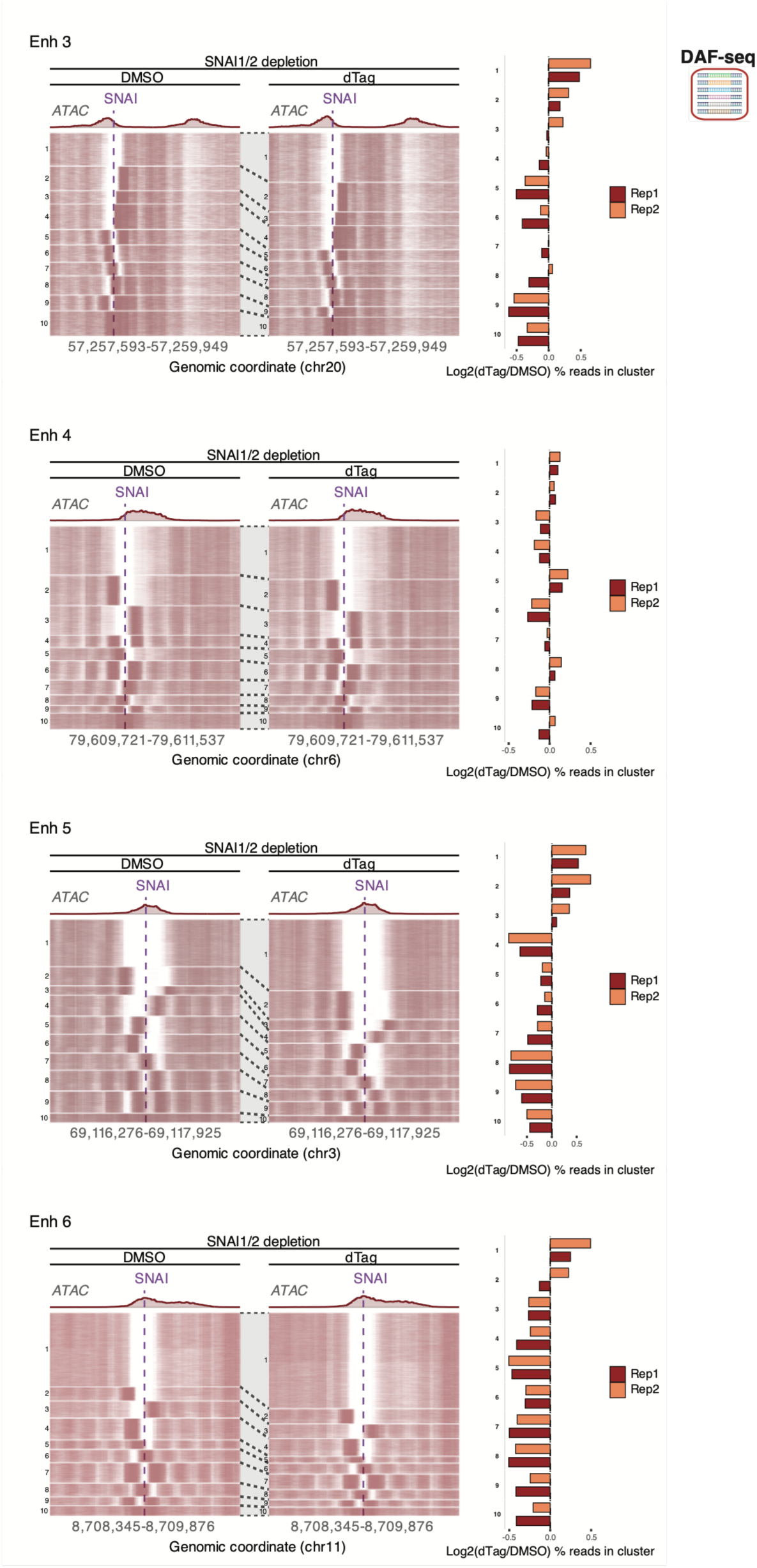
related to Figure 5: Single-fiber nucleosome states at tuned enhancers 3–6 upon SNAI1/2 depletion. Single-fiber nucleosome heatmaps for enhancers 3, 4, 5, and 6, showing DAF-seq data (two biological replicates per condition, SNAI1-FNV SNAI2-KO CNCCs, DMSO or 3 h dTAG treatment). Each row is one fiber; colored lines are nucleosome footprints (FiberHMM). Fibers were jointly k-means clustered (k = 10) on binary nucleosome occupancy within a region including 100 bp before and after the SNAI1/2 motif. ATAC-seq tracks are shown above the heatmap for reference (DMSO and dTAG scaled together). Each heatmap is accompanied by a bar plot showing log2 ratio of the percentage of fibers assigned to each cluster in dTAG versus DMSO, shown separately for each biological replicate.

**Supplementary Figure 5.**
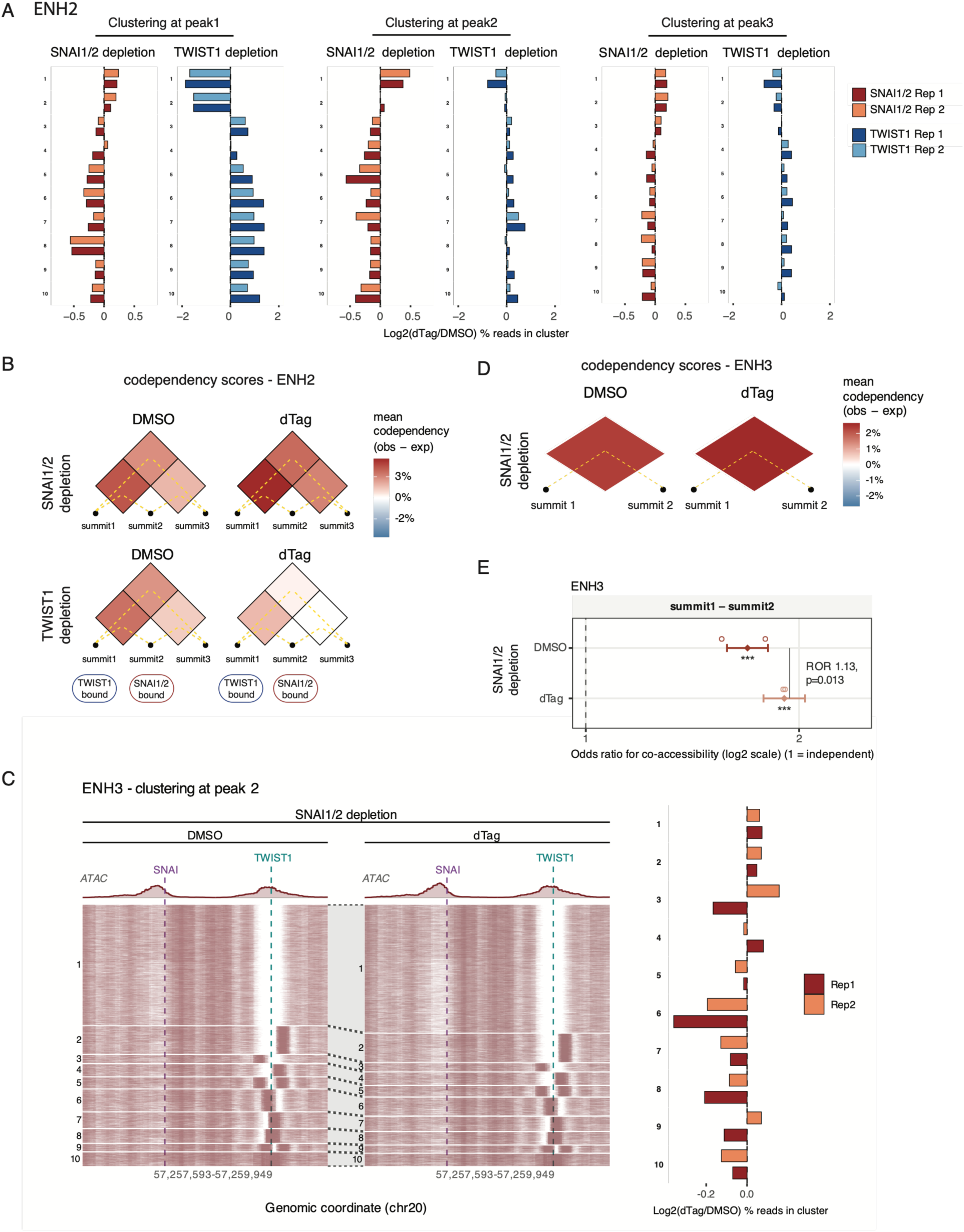
related to Figure 6: Chromatin-state changes and co-accessibility across peaks of clustered enhancers 2 and 3. (A) ENH2 chromatin states redistribution. Log2 ratio of the percentage of fibers assigned to each cluster (shown in figure 6B) in 3h dTAG versus DMSO, shown separately for each biological replicate. (B,D) LD-style triangle plots of fiber-level codependency calculated from DAF-seq data at the three ENH2 summits (B) or the two ENH3 summits (D). Fill = codependency, P(accessible at both) − P(acc1)×P(acc2), averaged over the two biological replicates: red, summits open together more often than expected under independence (cooperative); blue, less often (mutually exclusive); white, independent. All panels share one colour scale. A fiber was scored accessible at a summit if an MSP ≥100 bp overlapped it; marginals and the joint are computed on the same fibers, restricted to those spanning all summits. Odds ratios and pooled statistics for the same pairs are in Figure 6D and Table S4. (C) Single-fiber nucleosome heatmap for enhancers 3, clustered around the TWIST1 motif in peak 2. Shown is DAF-seq data (two biological replicates per condition, SNAI1-FNV SNAI2-KO CNCCs, DMSO or 3 h dTAG treatment). Each row is one fiber; colored lines are nucleosome footprints (FiberHMM). Fibers were jointly k-means clustered (k = 10) on binary nucleosome occupancy within a region including 100 bp before and after the TWIST1 motif in peak 2. ATAC-seq tracks are shown above the heatmap for reference (DMSO and dTAG scaled together). Each heatmap is accompanied by a bar plot showing log2 ratio of the percentage of fibers assigned to each cluster in dTAG versus DMSO, shown separately for each biological replicate. (E) Odds ratio for co-accessibility between pairs of ENH3 summits on individual DAF-seq fibers. A fiber was scored accessible at a summit if an MSP ≥100 bp overlapped it; only fibers spanning both summits were used. Filled diamonds, Mantel–Haenszel OR pooled across two biological replicates (stratified by replicate) with 95% CI; open circles, individual replicates (Fisher exact). Dashed line, OR = 1 (independence); OR > 1, summits open together more often than expected. x axis log2. Asterisks, BH-adjusted CMH p (* < 0.05, ** < 0.01, *** < 0.001). Brackets give the ratio of odds ratios (ROR = OR_dTAG/OR_DMSO) from a fiber-level logistic interaction model with replicate-specific intercepts, likelihood-ratio test, BH-adjusted.

**Supplementary Figure 6.**
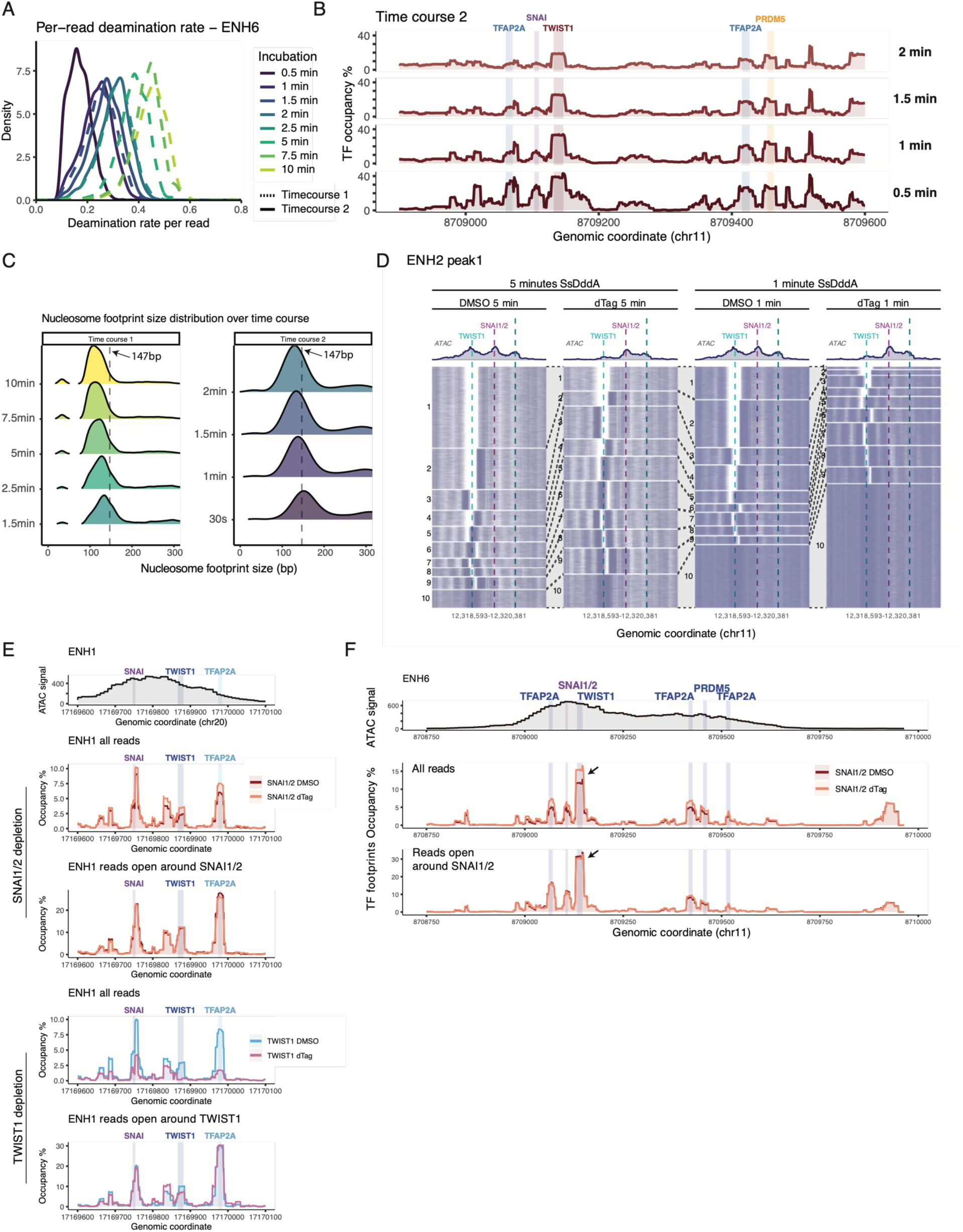
related to Figure 7: Short deaminase pulses resolve TF footprints without altering nucleosome state distributions. (A) Distribution of deamination rates per read (DAF-seq) comparing two independent experiments (time course 1 and 2) with different SsDddA incubation times (enhancer 6). (B) Metaplots of TF footprint occupancy percentage for enhancer 6, with the indicated SsDddA incubation times (time course 2 samples). An ATAC-seq track (DMSO) is shown above the metaplot for reference. (C) Distribution of nucleosome footprint sizes comparing different SsDddA incubation times. (D) Single-fiber nucleosome heatmaps for enhancer 2, clustered around TWIST1 motif in peak 1, comparing DMSO and dTAG samples of TWIST1-FV line treated with 1 minute or 5 minutes SsDddA. All fibers were clustered together. Each row is one fiber; colored lines are nucleosome footprints (FiberHMM). ATAC-seq tracks are shown above the heatmap for reference (DMSO and dTAG scaled together). (E-F) Metaplots of TF footprint occupancy percentages for enhancers 1 and 6, comparing DMSO and 3h dTAG conditions for either SNAI1-FNV SNAI2-KO or TWIST1-FV CNCCs. For each line, shown is a metaplot normalized over all reads and a metaplot of only reads belonging to clusters with extended nucleosome-free regions around the respective TF (ENH1: cluster 1 [Figure 5C], ENH6: cluster 1 [Figure S4]). An ATAC-seq track (DMSO) is shown above the metaplot for reference.

## Declaration of generative AI and AI-assisted technologies in the manuscript preparation process

During the preparation of this work, the authors used Claude (Anthropic) for drafting, reviewing and debugging the data-processing scripts described in methods and available at [https://github.com/luciaichino/SNAI1-2_CNCC], and to edit the language of the manuscript. The authors reviewed and edited the output as needed and take full responsibility for the content of the published article.

